# Genomic differentiation among European perch in the western Baltic Sea reflects colonisation history and local adaptation

**DOI:** 10.1101/2022.11.23.516742

**Authors:** Mikkel Skovrind, George Pacheco, Emil Aputsiaq Flindt Christensen, Shyam Gopalakrishnan, Katharina Fietz, Tore Hejl Holm-Hansen, Filipe Garrett Vieira, Marcus Anders Krag, Henrik Carl, M Thomas P Gilbert, Morten Tange Olsen, Peter Rask Møller

## Abstract

Environmental variation across the distribution of wild species can lead to local adaptations. The Baltic Sea was formed when the Fenno-Scandian ice sheet retreated around 12 thousand years ago, creating a new brackish water habitat colonised by both marine and freshwater fish species. The European perch (*Perca fluviatilis*) is a predatory freshwater fish with a large geographical distribution across Eurasia, where it inhabits a wide range of environmental niches. In the Baltic Sea region it has even developed a specialised brackish water phenotype that can tolerate environmental salinity levels, which are lethal to the ancestral freshwater phenotype. However, very little is known about the colonisation history and underlying genomic mechanisms facilitating the colonisation and adaptation of perch to the Baltic Sea. Here, we use Genotyping-By-Sequencing data from six freshwater and six brackish water localities to disclose the evolutionary relationship between the freshwater and brackish water phenotype. Our results show that the brackish water perch phenotype occurs in multiple distinct genetic clusters. We find that gene flow between brackish water phenotypes with full access to the sea likely led to lower levels of differentiation and higher diversity than in freshwater phenotypes. Selection analyses suggest that genomic adaptation played a role in the colonisation of the Baltic Sea and that the top three regions under selection harbour salinity tolerance genes. We also find a link between the historic salinity of the Baltic Sea and the demographic history of the brackish water phenotypes and go on to discuss the implications of our findings for management of brackish water perch in the western Baltic sea.

**Highlights:**

- GBS data from 12 perch populations, six with brackish and six with freshwater origin
- Colonisation history and differentiated gene flow shaped the current population structure
- The brackish water ecotype was found in all three major genetic clades
- Top three regions under selection harboured salinity tolerance genes
- Salinity influenced Ne during the formation of the Baltic Sea

**Graphical abstract:** 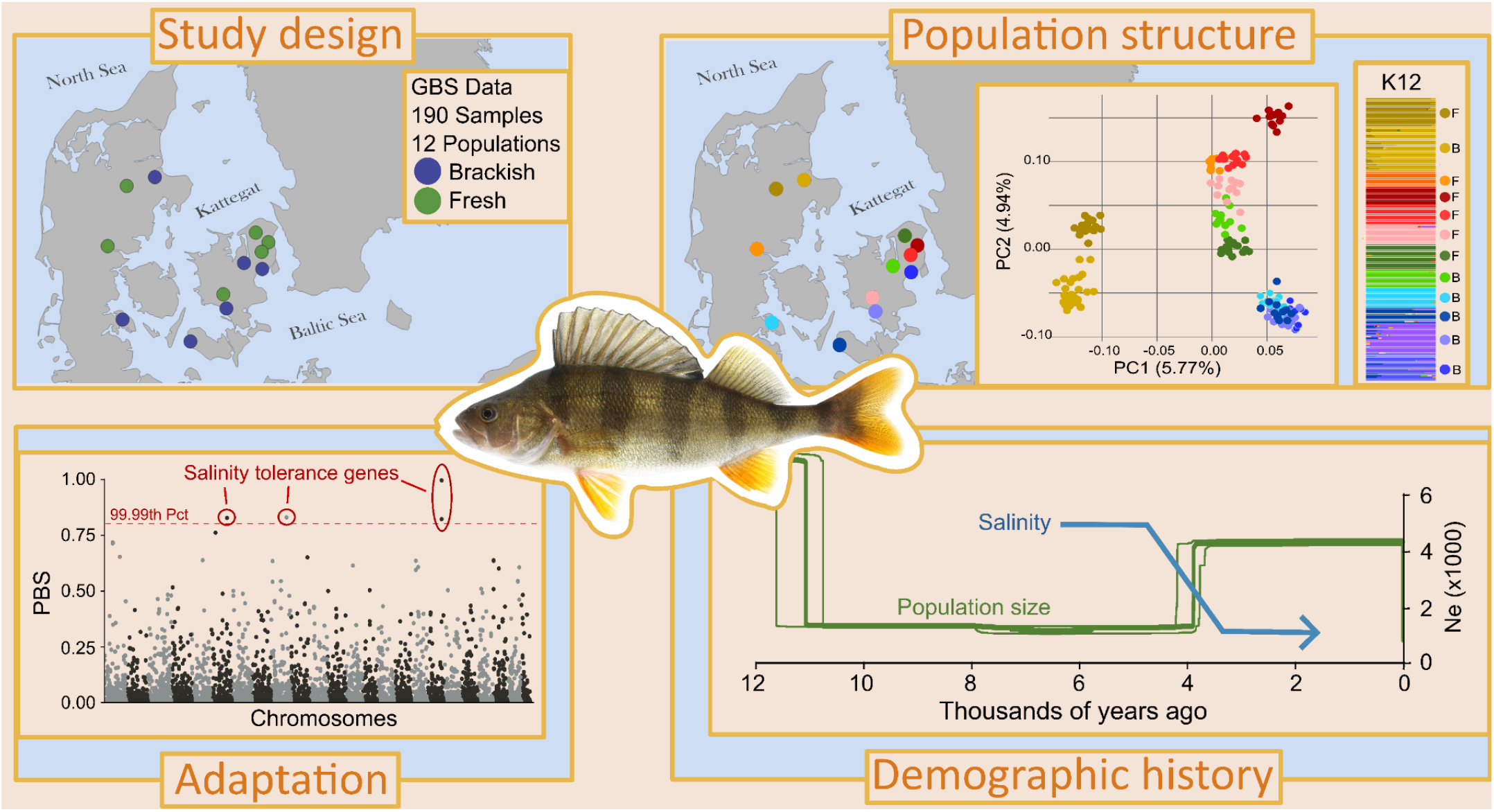

## Introduction

Environmental heterogeneity plays a key role in shaping intraspecific variation in wild species (McDonald & Ayala, 1974). Novel habitats can offer new opportunities and challenges to populations in a subsection of the species distribution range, thus promoting local adaptation and increased levels of differentiation (Langerhans et al., 2013). Hence, intraspecific variation in morphology, behaviour and physiological capabilities is often correlated with environmental variation and can over time lead to the formation of separate ecotypes (Foote et al., 2016; Skovrind et al., 2016). While phenotypic plasticity can be a vector towards adaptation (Radersma et al., 2020), genomic diversification is often the key mechanism governing the formation of ecotypes (Feder et al., 2012; Foote et al., 2016; Seehausen et al., 2014). When populations inhabit separate niches and no longer interbreed, genetic drift and/or selection will affect them separately and increase the differentiation between them across the genome and in particular in genes underpinning local adaptation.

Almost all teleost species have internal osmotic pressures corresponding to salinities of 9 - 12 parts per thousand (ppt) (Brett, 1979), but have evolved to live in the stable salinity environment of either freshwater (around 0 ppt) or marine habitats (around 35 ppt). Even subtle variations in internal osmotic pressure can compromise their physiological well-being and be lethal (Brauner et al., 1992; Christensen et al., 2019; Lutz, 1972). To counterbalance adverse effects of alternated internal salinity, fish osmoregulate (E. H. Larsen et al., 2014). Although fish are the most diverse vertebrate taxa in the world (Magurran et al., 2011), relatively few species are capable of both osmoregulating at ambient salinities below their internal osmotic pressure and ambient salinities above their internal osmotic pressure (Evans, 1984). Hence, teleost fish species diversity is generally low in intermediate and fluctuating salinities found in brackish water estuaries, compared to adjacent fresh and marine habitats (Remane, 1934; Whitfield et al., 2012). However, since the biological productivity of estuaries is high, species and specialised phenotypes that adapt to this challenging environment can benefit from high food availability and low interspecific competition.

The Baltic Sea is the world’s largest estuary at 400,000 km^2^ and characterised by a marked salinity gradient from nearly freshwater (<0.5 ppt) in the north to fully marine (35 ppt) as it merges with the belts and straits (35 ppt) in the west (Leppäranta & Myrberg, 2009; Weckström et al., 2017). In the southwestern Baltic Sea, the environmental salinity fluctuates between 7 and 22 ppt making it a challenging habitat requiring both osmoregulation below and above iso-osmotic levels (hyper and hypo-osmoregulation, respectively) (Weckström et al., 2017). In this region, a few species of fish adapted to freshwater (stenohaline freshwater fish) have brackish phenotypes utilising the greater resources in the brackish environment, e.g. ide (*Leuciscus idus*) (Skovrind et al., 2016), northern pike (*Esox lucius*) (Jacobsen et al., 2017) and European perch (*Perca fluviatilis*)(Christensen et al., 2021; Skovrind et al., 2013). Stenohaline freshwater fish are adapted to ambient salinities lower than their internal osmotic pressure, and hyper-osmoregulate by secreting diluted urine and taking up specific ions from the environment (E. H. Larsen et al., 2014). Interestingly, though, in the intermediate salinities of the southwestern Baltic Sea, populations of stenohaline freshwater fish have developed the ability to hypo-osmoregulate at salinities higher than their internal osmotic pressure, which requires fundamentally different osmoregulation mechanisms involving gastro-intestinal uptake of imbibed ambient water, and excretion of excess ions (E. H. Larsen et al., 2014). These enhanced osmoregulation capabilities of brackish water phenotypes of stenohaline freshwater fish makes them ideal models for studying intraspecific variation between phenotypes and adaptation to a novel environment.

Through the analyses of Genotype-By-Sequencing (GBS) (Elshire et al., 2011) data of 190 individuals from six brackish water and six freshwater perch populations in the southwestern Baltic Sea region (Figure 1), we provide novel insights into the evolution of the two phenotypes. Specifically, we infer the genomic population structure of these populations and identify genomic regions under selection in the brackish water phenotype, which could explain the increased ability to thrive in brackish water. We also estimate the demographic history of the brackish water phenotype during the historic changes in environmental salinity in the Baltic Sea.

**Figure 1.**
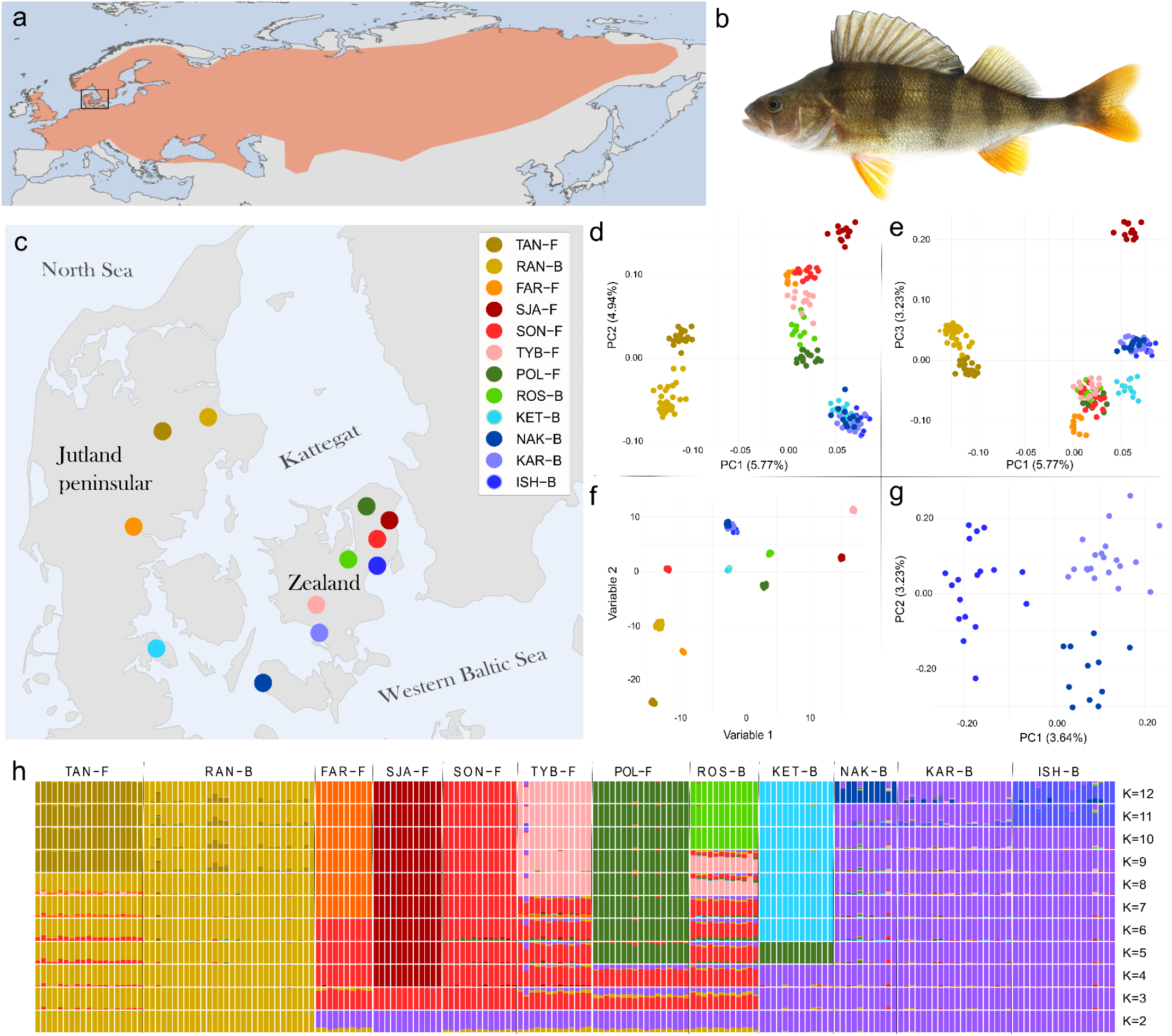
Population structure of perch in the southwestern Baltic Sea region. **a)** Native distribution of perch *Perca fluviatilis*. **b)** A 24.5cm European perch from Køge harbour on the east coast of Zealand December 20th 2022 (photo by Henrik Carl). **c)** Brackish water and freshwater sample sites included in the present study of Baltic Sea perch. -B indicates brackish water and -F indicates freshwater localities. **d-g)** Principal Component Analyses. Panels **d** and **e** are the PCAngsd results for PC1-PC2 and PC1-PC3, respectively. Panel **f** is the UMAP analyses, which used the PCAngsd results as input. **g)** Principal Component Analysis of brackish water perch from NAK-B, KAR-B and ISH-B. **h)** Ancestry proportions estimated with NGSadmix. Vertical bars represent single individuals. Different colours indicate the estimated ancestry proportions in each individual.

## Methods

### Sampling

A total of 190 fin-clip samples were collected between September 2013 and April 2014 at 12 localities (Figure 1b); six freshwater localities and six brackish water localities (Table 1). The brackish water localities, which all harbour anadromous perch, fell into two categories; (i) perch with full access to the brackish water sea (NAK-B, KAR-B and ISH-B); and (ii) perch inhabiting a brackish water fjord, but isolated from other brackish water perch populations by high saline marine water (RAN-B, ROS-B and KET-B). The number of samples per population ranged from 10 to 30. All samples were stored in 96-99 % ethanol at -18°C. The fish were individually numbered and adult individuals were preserved at the National History Museum of Denmark (See Table S1 for ID numbers). Permission for scientific fishing was provided by the Danish Ministry for Food, Agriculture and Fishery (journal no. 2009-02530-23088).

### Data generation

Genomic DNA extractions were performed using the DNeasy Blood & Tissue Kit (Qiagen, Valencia, CA) following the manufacturer’s protocol. The extracts were quantified using the Qubit 2.0 Fluorometer (Thermo Fisher Scientific, Waltham, MA). To check for molecular integrity, an aliquot of each DNA extract was run on a 1% agarose gel against a 1-kb ladder. We sent 190 extracts that passed our filters (minimum DNA concentration of 10 ng/μL and average fragment size above 20 kb) to the Institute for Genomics Diversity at Cornell University, where the GBS method was applied (Elshire et al., 2011). Samples were sent in two 96-well plates, each including a negative control. Once at Cornell, the DNA extracts were treated with the restriction enzyme EcoT22I before library preparation. All libraries had appropriate concentration, fragment size distribution and minimal adapter dimers, thus passing quality control. The libraries of each plate were pooled separately and sequenced on the HiSeq 2000 (Illumina, San Diego, CA) using a single-end 100 bp technology.

### GBS data processing, filtering and mapping

The raw sequencing GBS data was demultiplexed using GBSX v1.3 (Herten et al. 2015) allowing for one mismatch in the barcodes and one mismatch in the enzyme cut-site while retaining common sequencing adapters. Chimeric GBS reads were identified as discussed in Pacheco et al. (2020). In short, these reads occur when reads from two or more biological cut-sites are merged into a single artificial read. Thus, chimeric reads were defined as reads with more than one cut-site that mapped to two or more noncontiguous regions in the reference genome. Chimeric reads could bias our coverage statistics, so they were excluded using a script available at: https://github.com/g-pacheco/PerchGenomics.

The filtered sequencing data was mapped to the GENO_Pfluv_1.0 European perch reference genome (NCBI accession: GCA_010015445.1). The mapping was restricted to the 24 nuclear chromosomes thus excluding additional unplaced scaffolds. Analyses were further restricted to the fraction of the genome theoretically available to the GBS method. To determine this fraction, we performed an in-silico digestion on the reference assembly with the same enzyme used in the GBS protocol (EcoT22I) using BioSeq v/1.11 (Cock et al., 2009), and considered only the regions spanning 93 bp downstream and upstream each locus. Loci located less than 93 bp apart from each other, were merged into single-locus loci. All regions of interest were listed in a bed file available from the GitHub data repository (see link in the data availability section).

Samples were excluded if they had low amounts of data. To identify these samples, we created a presence/absence matrix for all loci, where loci covered by a minimum of three reads were scored as present and loci covered by fewer than three reads were scored as absence. Due to the magnitude of the matrix, we clustered the loci (k-means with K = 300 clusters). The resulting matrix was plotted as a heatmap with the samples hierarchically clustered by employing the R package pheatmap (Kolde, 2012). This heatmap (Figure S1) was visually inspected, and samples that clustered with the negative controls were excluded from further analyses.

The data was further processed and filtered using ANGSD v0.935 (Korneliussen et al., 2014). Aiming to remove poor quality data, we removed reads with a flag above 255, removed reads with multiple best hits and performed standard BAQ computation. We also adjusted mapping qualities for excessive mismatches to the reference, ignored bases with base qualities below 20, and removed reads with mapping qualities below 30. We excluded sites with more than five percent missing data across all samples. Sites with less than three reads per individual were called as missing data. We calculated genotype likelihoods using the SAMtools approach(H. Li et al., 2009), and sites were only identified as variable if they had a p-value below 1e-6, according to a likelihood ratio test, using a chi-square distribution with one degree of freedom. Sites with excess coverage were removed as they could belong to paralogous regions. To identify these sites we used ANGSD to calculate the sequencing depth and created a density plot using ggplot2 (Wickham, 2016). After visually inspecting the global depth plot (Figure S2), we set the maximum global depth at 190 times the number of individuals. Unless otherwise mentioned all subsequent analyses were performed using these filters.

### Population genetic structure

The population genetic clustering of samples was estimated with PCAngsd (Meisner & Albrechtsen, 2018) applying a minimum minor allele frequency of 0.006, corresponding to two copies of the minor allele in the dataset. Traditional principal component (PC) plots were made of PC1 against PC2 and PC1 against PC3. However, the covariance matrix generated by PCAngsd included ‘*n’* number of dimensions (in our case 188). In order to evaluate the information stored in all these dimensions, we applied the dimensions reduction software UMAP (Becht et al., 2018), which reduced the complexity of our 188-by-188 matrix to two dimensions making it possible to capture and visualise all dimensions in a single plot. To further investigate the relationship between closely related populations, we also performed a PCAngsd analyses only including the brackish water perch from NAK-B, KAR-B and ISH-B, using the settings described above, except the minimum minor allele frequency, which was set to 0,025, again corresponding to two alleles. To further estimate genetic clusters based on individual admixture proportions we used NGSadmix v3.2 (Skotte et al., 2013) for 2-12 ancestral populations (K-values), using default parameters, except for tolerance for convergence which was set to 1e-6, log likelihood difference in 50 iterations, which was set to 1e-3 and maximum EM iterations which was set to 10,000. For each K-value, 100 replicates were applied and the replicate with the highest likelihood was used for subsequent plotting and interpretation.

### Diversity, differentiation, geographic distance and salinity

ANGSD was used to compute the unfolded site allele frequency (SAF) of each population using the closely related North-american sister species, the yellow perch *Perca flavescens* (accession number: SAMN10722690) as ancestral state. The SAF was subsequently used to estimate the Site Frequency Spectrum (SFS) for each population and the two dimensional SFS (2dSFS) for each population pair using realSFS (Mas-Sandoval et al., 2022). The SFS was used to estimate the nucleotide diversity (*π*) of each population and the 2dSFS was used to estimate the pairwise differentiation (*F*_ST_) among populations. To estimate the relation between geographic distance and genomic diversification (Wright, 1943), we plotted the *F*_ST_ values against the euclidean distance, measured using Google Earth (available from https://earth.google.com/) of all sample site pairs. Correlations were assessed for all sample sites, as well as brackish water pairs and freshwater pairs separately using a linear model. To assess the effect of salinity on perch genetic variation, including the salinity levels that perch migrating to and from brackish water sample sites would have to endure, environmental salinity data for the sea immediately adjacent to the fjords or estuaries associated with the brackish water sample sites were retrieved. All salinities were extracted once daily for the period 01-09-2013 to 31-08-2014 from the models created by MyOcean (accessible at: www.myocean.eu). The mean annual salinity was then plotted against the nucleotide diversity to estimate the effect of isolation on genetic diversity.

### Local adaptation in brackish water perch

To identify regions of the genome showing signs of selection in the brackish water perch populations, which have increased salinity tolerance (Christensen et al., 2019), we calculated the Population Branch Statistics (PBS) in ANGSD (Korneliussen et al., 2014). PBS is a summary statistic that quantifies the genetic drift in one population, relative to two other populations, across the genome using a sliding window approach. The length of each branch of their corresponding three-population tree is estimated and windows with significantly longer branches indicate positive selection in the corresponding population. To increase the sample size, the PBS analysis was based on core populations belonging to the main genetic clusters that we identified (see results); Cluster A (RAN-B and TAN-F), Cluster B (SON-F and TYB-F), and Cluster C (NAK-B, KAR-B and ISH-B). The SFS was calculated for each of the groups using the same parameters used to calculate the summary statistics, with 50% missing data. The analysis was run with 50kb overlapping windows in 25kb steps. The generation time was set to 3.4 years, based on the median generation time of eight perch populations each estimated from the von Bertalanffy growth parameters published by Christensen et al. (2021) using the generation time estimation approach described in (Froese & Binohlan, 2000). The eight populations included five with freshwater and three with brackish water origin; parameter details can be found in Table S2. As mutation rates are unavailable for the *Perca* genus, we applied a per generation mutation rate of 3.7×10-8 estimated for three spined sticklebacks (*Gasterosteus aculeatus*) (S. Liu et al., 2016). To identify genes and their functions in the genomic regions most likely to be under positive selection, we blasted the annotated protein sequences from the reference genome using BlastP (Johnson et al., 2008). We included protein sequences from genes 100kb either side of each window and used the result to search the Gene Ontology AmiGO2 database (Ashburner et al., 2000; Carbon et al., 2009; The Gene Ontology Consortium, 2021) for the corresponding genes and their molecular functions. Subsequently, literature searches were performed for all gene names in association with descriptive phrases “salinity tolerance”, “salt tolerance” and “osmotic stress”.

### Demographic history

To estimate recent demographic changes in the brackish water perch in the southwestern Baltic region, we ran StairwayPlot v2.1 (X. Liu & Fu, 2020) on the core populations of Cluster C (NAK-B, KAR-B and ISK-B) following the best-practice instructions available at: https://github.com/xiaoming-liu/stairway-plot-v2. We used the dataset with 50% missing data for the demographic analysis in order to have a sufficient amount of variable sites and cover a larger proportion of the genome. We applied a generation time of 3.4 years and mutation rate of 3.7×10-8 as described above. The usefulness of GBS data for demographic inferences with the Stairway Plot method was further demonstrated by (Hansen et al., 2018) who trimmed and downsampled high coverage whole genome sequencing data to mimic GBS reads and were able to recapture the demographic trajectory of the original data. This gives us confidence that our demographic results are reliable.

## Results

### Data summary

Each sample locality included between 10 and 30 samples (mean = 15.8). Locality details can be found in Table 1 and sample details can be found in Table S1. After inspecting the presence/absence heatmap (Supplementary Fig. 2), the two blanks and two samples (NAK-B_09 and POL-F_15) that clustered with the control samples were excluded from further analyses.

### Population genetic structure

The PCA analyses separated perch by geography and salinity. PC1 (5.77% variation) separated the two localities in northern Jutland from the remaining localities, while PC2 (4.97% variation) separated the brackish water perch from the freshwater perch (Figure 1d). PC3 (3.23% variation) separated the KET-B locality from the other brackish water localities in western Baltic Sea (NAK-B, KAR-B and ISH-B) and separated the freshwater locality SJA-F from all other localities (Figure 1e). When applying the UMAP dimension reduction method to the PCA data, all sample sites formed their own non-overlapping clusters, except for the three brackish water localities in southeastern Zealand. The analysis, which only included these three localities (NAK-B, KAR-B and ISH-B) separated ISH-B from NAK-B, KAR-B on PC1, while NAK-B, KAR-B separated from each other on PC2 (Figure 1g). Thus, each of these three brackish water localities in southeastern Zealand formed their own genetic clusters at a finer scale.

The individual admixture coefficients estimated for K=2 to K=12 supported the PCA and revealed further fine-scale population genetic structure among perch localities in the western Baltic Sea region (Figure 1h). In the analyses of two ancestral populations (K=2) the two most northern localities on the Jutland peninsula (TAN-F and RAN-B) formed a separate cluster (Cluster A). At K=3, the remaining populations were separated into two clusters, one (Cluster B), which included the remaining freshwater localities (FAR-F, SJA-F, SON-F, TYB-F, POL-F, and a single brackish water locality ROS-B), and the other (Cluster C), which included the remaining brackish water localities (KET-B, NAK-B, KAR-B and ISH-B). For K=4 to K=12, additional genetic structuring was detected with all 12 localities comprising their own genetic cluster at K=12. Admixture (gene flow) was detected from TAN-F to RAN-B in Jutland and among the three brackish water populations NAK-B, KAR-B and ISH-B in southeastern Zealand.

### Diversity, differentiation, geographic distance and salinity

The highest levels of nucleotide diversity (*π*) was found in the brackish water localities NAK-B, KAR-B and ISK-B, which had *π* estimates between 0.000360 and 0.000370 (Figure 2). Intermediate levels of diversity was observed in TYB-F and ROS-B with *π* estimates between 0.000339 and 0.000345 while the remaining localities (TAN-F, RAN-B, FAR-F, SJA-F, SON-F, POL-F and KET-B) had *π* estimates below 0.000307. The lowest *F*_ST_ values, which were found among the southeastern brackish water localities NAK-B, KAR-B and ISH-B, all ranged from 0.032 to 0.029. All other *F*_ST_ values ranged between 0.100 and 0.278 with the highest values being found between FAR-F and KET-B. The IBD analyses revealed that there was very little correlation between genetic diversity (*F*_ST_) and geographic distance (Km) between localities (figure S5). The weakest correlation was found when including only the freshwater localities (R^2^=0.002), and the strongest correlation was found when including only the brackish water localities (R^2^=0.308). The analysis of all localities revealed intermediate correlation (R^2^=0.108). The environmental salinity data showed that perch migrating to and from the brackish water localities NAK-B, KAR-B and ISH-B would most of the year experience salinity levels lower than the 18 ppt physiological limit of perch, implying that immigration and emigration at these localities is not limited by salinity (Figure 3a-b). In contrast, migrants to and from the KET-B and ROS-B localities would experience salinities above the physiological limits approximately half the days with ROS-B periodically experiencing much higher salinity levels. The perch migrating to and from the RAN-B locality would experience higher salinities than any other locality with more than half the observed days above 19.5 ppt. Overall, our results further showed that there was a strong negative correlation (R^2^=0.7132) between genetic diversity (*π*) and the salinity levels any putative migrants would have to endure (Figure 3c).

**Figure 2.**
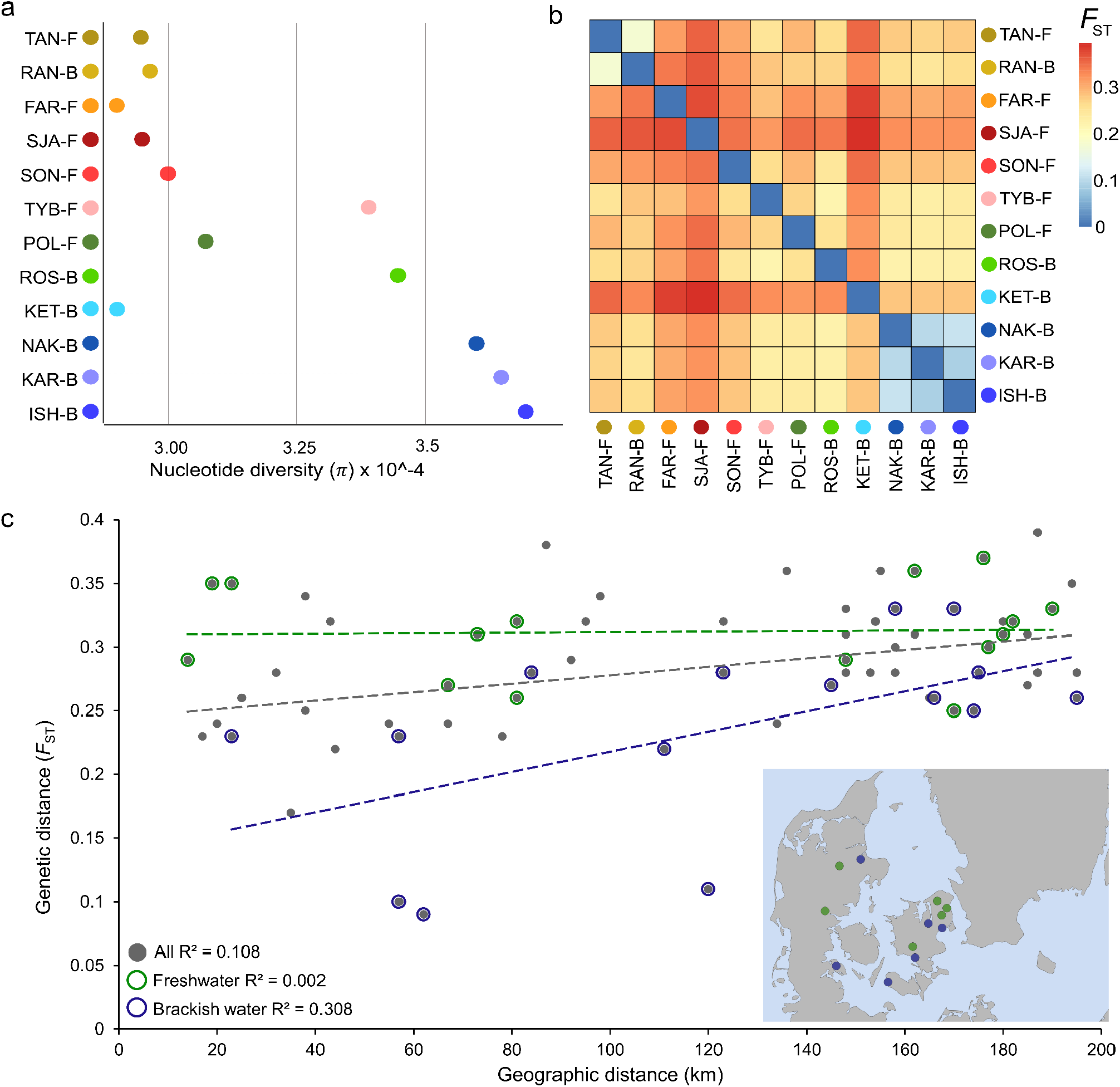
Diversity and differentiation of perch in the western Baltic Sea region. **a)** Genomic nucleotide diversity (π). See Figure 1 for locality names and geographic placement. -B=brackish; -F=freshwater. **b)** Population differentiation heatmap based on *F*_ST_ values. **c)** Isolation by distance. Linear correlations between genetic (*F*_ST_) and geographic (Km) distance among perch populations in the western Baltic Sea region.

**Figure 3.**
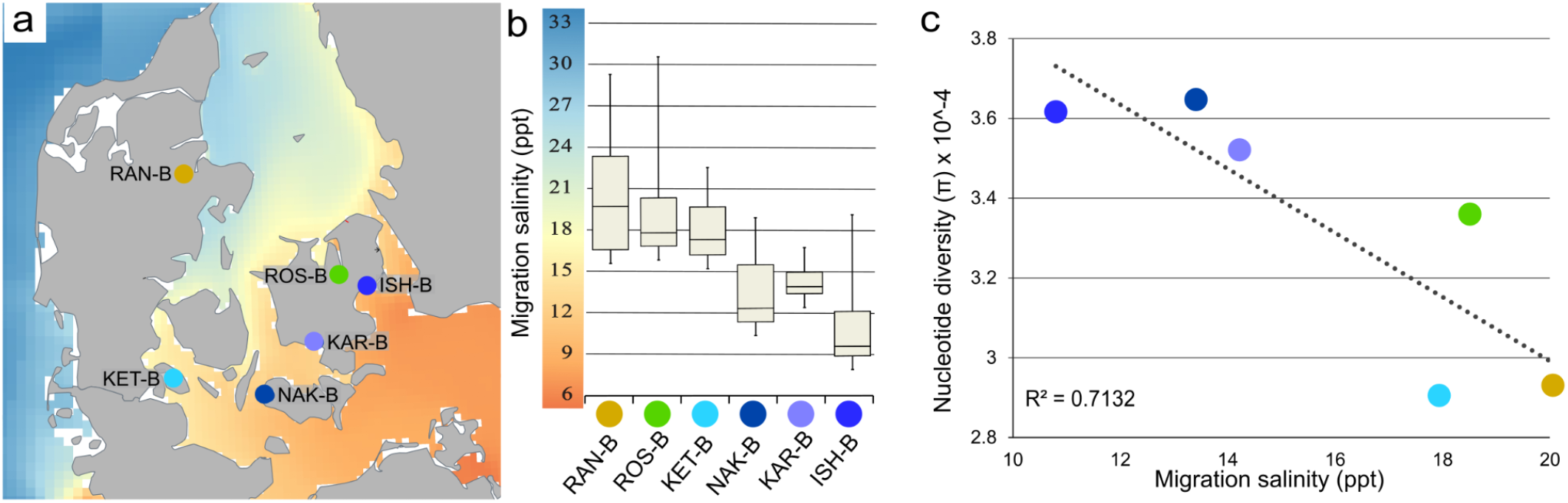
Modelled environmental salinity of the western Baltic Sea and correlation with genetic diversity. **a)** Average annual environmental salinity based on one daily data point spanning from September 1st 2013 to August 30th 2014. **b)** Environmental salinity outside the stream/river/fjord associated with each brackish water population. **c)** Correlation between genetic diversity and salinity.

### Local adaptation in brackish water perch

In order to identify genomic regions under selection in brackish water perch, PBS analyses were performed on 26,165 windows with an average of 278 nucleotide sites per window. The average genome wide PBS values for Clusters A, B and C were 0.019, 0.024 and 0.016, respectively. In Cluster C, comprised of the four brackish water localities (KET-B, NAK-B, KAR-B and ISH-B), the three genomic regions with the highest PBS values, which were all above 0.82, were found in chromosome 6 (range: 30650K-30925K), chromosome 9 (range: 32550K-32825K) and chromosome 18 (range: 28775K-29050K), each including seven, six and 12 protein coding genes, respectively (Figure 4a; Table S3; Table S4).

**Figure 4.**
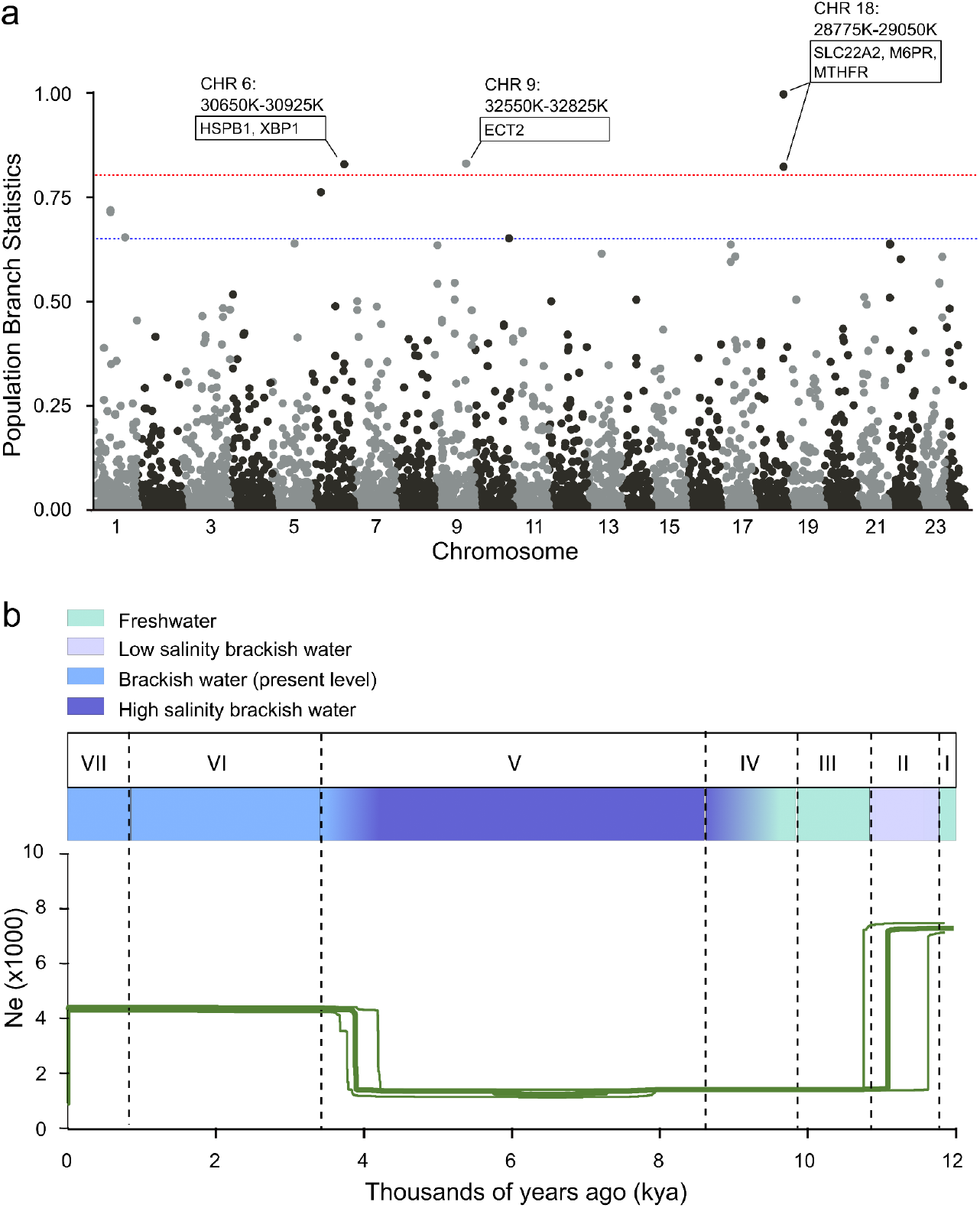
Adaptation and demographic history of brackish water perch in the western Baltic Sea (NAK-B, KAR-B and ISH-B). **a)** Identification of regions and genes under selection using Population Branch Statistics (PBS) performed with 50kb windows in 25kb steps. Grey and black points indicate results from uneven and even chromosome numbers, respectively. The blue and red horizontal lines represent the 99.5th and 99.9th percentile, respectively. **b)** Demographic history of brackish water perch from the western Baltic Sea. I-VII indicates the environmental phases of the Baltic Sea (Phase I: Baltic Ice Lake, Phase II: Yolandia Sea, Phase III: Ancylus Lake, Phase IV: Initial Littorina Sea, Phase V: Littorina Sea, Phase VI: Post-Littorina Sea, Phase VII: Modern conditions (Weckström et al., 2017).

### Demographic history

The demographic history of the brackish water perch at NAK-B, KAR-B and ISH-B showed a decline in effective population size coinciding with the formation of the Baltic Sea 11-12 kya from 7000 to 1000 individuals (Figure 4b). The effective population size was stable at this level until ∼4 kya when it increased to 4400 individuals, coinciding with a period of Baltic Sea environmental changes from high salinity brackish water to lower contemporary salinity levels. Runs excluding singletons gave near identical results (not shown).

## Discussion

### Patterns of genetic variation in the western Baltic Sea perch

Unravelling a species’ population structure is a first step in attempting to understand the evolutionary processes that shaped the present day genetic variation among wild populations. Our data revealed that each sample locality constituted a separate genetic population and there was no evidence of substructure within populations. Isolation by geographical distance played a limited role in the genetic differentiation of populations. Rather, we suggest that colonisation history and substantial salinity gradients have been the main drivers of differentiation among perch populations in the region. Specifically, all analyses found localities in northern Jutland to form a separate genetic cluster (Cluster A) and be highly distinct from all other perch localities in our study. This deep genetic split likely represents colonisation from two different refugia after the retreat of the Fenno-Scandian ice sheet at the Pleistocene-Holocene transition. The existence of multiple past glacial refugia has previously been suggested as the underlying mechanism behind large scale population structure in both European perch (Christensen et al., 2016; Nesbø et al., 1999; Toomey et al., 2020) and other freshwater fish species in northern Europe (Bekkevold et al., 2015; Culling et al., 2006; Kontula & Väinölä, 2001; Skovrind et al., 2016). The other localities were split into two clusters; the first (Cluster B) comprised mainly of freshwater populations, as well as a single isolated brackish water population (ROS-B), and the second (Cluster C) including all southeastern brackish water localities with access to the western Baltic Sea.

Freshwater fish populations are usually separated in lakes or drainages with very limited opportunities for gene flow, however the colonisation of the brackish water environment can enable migration, thus facilitating gene flow (Skovrind et al., 2016). In our study, the freshwater localities and isolated brackish water localities in Clusters A and B followed a characteristic genetic freshwater pattern (Ward et al., 1994), with lower levels of diversity and higher levels of differentiation. This pattern is most likely the result of genetic drift acting on smaller and/or isolated populations, which have had limited contact with other populations since the colonisation (Nei & Tajima, 1981). The population structure in Cluster C composed of brackish localities followed a different pattern, most likely driven by differences in environmental salinity. The southeastern brackish water localities (NAK-B, KAR-B and ISH-B) were closely related and had high levels of diversity, while KET-B was more distantly related and had a low level of diversity. This could be because the sea surrounding the southeastern brackish water localities have environmental salinity within the tolerance of brackish water perch, thus allowing them to fully exploit the resources of the highly productive Baltic Sea and allow gene flow among them, while gene flow to and from KET-B was likely more restricted due to higher salinity levels, and perhaps deep waters (see e.g. Olsson et al. (2011)).

An influx of migrants bringing novel alleles into or among the populations would also explain the elevated levels of diversity, observed in NAK-B, KAR-B and ISH-B. The NGSadmix analyses suggest that admixed individuals are found in these three populations, supporting that gene flow plays a role in the high levels of diversity and lower differentiation. Another possible explanation for the high levels of diversity is that access to large marine habitats has allowed the western Baltic Sea brackish water populations to maintain large populations, which would minimise the loss of diversity to genetic drift. The very close relatedness of NAK-B, KAR-B and ISH-B could also indicate recent divergence. However, high diversity and presence of recently admixed individuals suggests that gene flow is at least partly behind the observed population structure pattern in Cluster C.

Intriguingly, while NAK-B, KAR-B and ISH-B were grouped with the same genetic cluster and likely had substantial gene flow, we did detect substructure among them, suggesting that these southwestern Baltic brackish water perch exhibits site fidelity to their natal stream. This is in accordance with previous studies that found genetic differentiation among perch populations in the inner Baltic Sea sampled less than one kilometre apart (Bergek & Björklund, 2009), as well as genetic differentiation within a single lake (Bergek & Björklund, 2007). Other mechanisms maintaining the differentiation among perch populations have previously been suggested, including lower fitness of F1 hybrids (Behrmann-Godel & Gerlach, 2008) and a preference to their own population based on olfactory cues (Behrmann-Godel et al., 2006).

### Local adaptation in brackish water perch

Our results suggest that selection acted on genomic variation and possibly facilitated the colonisation of the fluctuating salinity brackish waters of the southwestern and western Baltic Sea. The three genomic regions with the strongest signals of selection all included genes linked to salinity tolerance in other organisms. The strongest signal of selection was found in chromosome 18 where the analysis identified three genes linked to salinity regulation; SLC22A2, MTHFR and IGF2/M6PR. The SLC22A2 gene encodes a protein that transports organic cations across basolateral membranes. SLC22A2 has previously been shown to be differentially expressed in Pacific spiny dogfish (*Squalus suckelyi*) exposed to different salinity levels (Cole, 2018) and organic cation transporter genes has previously been identified as under selection in brackish water pike in the Baltic Sea (Sunde et al., 2022). The MTHFR gene encodes a rate-limiting enzyme in the methyl cycle and is crucial for the formation of methionine (vitamin B9). A study of three-spined sticklebacks (*Gasterosteus aculeatus*) showed that MTHFR was expressed at significantly different levels in saltwater and freshwater exposed individuals. Vitamin B9 has even been demonstrated to mitigate the effect of elevated salinity in barley (*Hordeum vulgare*) (Özmen & Tabur, 2020). The IGF2/M6PR gene encodes a transmembrane protein (El-Shewy & Luttrell, 2009), which has been shown to be differentially expressed under different salinity regimes in several fish species, including Arctic charr (*Salvelinus alpinus*) and half smooth tongue sole (*Cynoglossus semilaevis*) (S. Li et al., 2020; Norman et al., 2011). The second strongest signal of selection was located in a region of chromosome 9, which included the ECT2 gene, playing a vital role in stabilising mRNA in the cytoplasm and bindinding to and promotes the transcription of N6-methyladenosine in a wide array of plant species subject to salinity stress, including cotton (*Gossypium* sp.), sweet sorghum (*Sorghum bicolor)*, sugar beet (*Beta vulgaris)* and arabidopsis (*Arabidopsis* sp.) (Cui et al., 2022; Wang et al., 2022; Zheng et al., 2021). The third strongest signal of selection was identified on chromosome 6 and included three salinity tolerance associated genes (XBP1 and the two heat shock proteins HSPB8 and HSP67BA). The XBP1 gene has been shown to be upregulated in red-eared slider turtles (*Trachemys scripta elegans*) exposed to 15 ppt salt water and suggested as a driver of adaptation to brackish water (N. Li et al., 2021). Heat shock proteins are involved in the response to osmotic stress (Deane & Woo, 2011; Sørensen et al., 2003) and have been shown to be expressed differently in European flounder (*Platichthyes flesus)* translocated from marine to brackish water (P. F. Larsen et al., 2008). However, the resolution of GBS data is limited as it only captures genomic regions adjacent to cut sites and we are therefore not able to identify the specific genomic sites under selection, but only the general region. Thus, the signal of selection identified in our data is likely due to linkage between the GBS sites and the actual site or sites under selection.

Previous studies have shown that the brackish water perch phenotype has a markedly increased salinity tolerance of at least 17.5 ppt achieved through an ability to hypo-osmoregulate that freshwater conspecifics have not (Christensen et al., 2017, 2019). The identification of salinity tolerance associated genes within each of the three genomic regions with the strongest signals of selection suggest that increased physiological abilities to cope with saline environments of the brackish water phenotype is rooted in genomic adaptation and not a result of phenotypic plasticity alone. Combined with the physiological and behavioural (anadromous versus resident) differentiation between brackish and freshwater perch, our results suggest that the brackish water perch should be considered a separate ecotype. The Baltic Sea was formed relatively recently in evolutionary time, ∼12 thousand years ago after the Pleistocene/Holocene transition when the Fenno-Scandian ice sheet retreated (Björck, 1995; Hall & van Boeckel, 2020). Thus, the adaptation to hypo-osmoregulation observed in our data likely arose within this period. Such rapid evolution has also been described in three-spined sticklebacks, which have adapted in parallel to freshwater habitats upon post-glacial isolation from marine environments (Hohenlohe et al., 2010). Our limited sample sizes only allowed for analyses of selection in the southeastern brackish water localities, but it is likely that similar adaptations occurred in parallel in the isolated brackish water populations KET-B, RAN-B and ROS-B and elsewhere. Indeed, the presence of brackish water phenotypes in all three main genomic clusters (i.e. Cluster A, B and C) suggest that adaptations to brackish water arose in parallel in different post-glacial colonisation waves. Future studies should seek to explore patterns of salinity tolerance and adaptations across multiple brackish and freshwater perch populations.

### Demographic history

Understanding the effect of environmental changes on population sizes through time are vital when trying to predict the impact of future changes. Our stairway plot analysis suggests an association between the historic environmental salinity of the western Baltic Sea and the demographic history of brackish water perch in the region. During the Baltic Ice Lake period (phase I in Figure 5), which followed the retrieval of the Fenno-Scandian ice sheet, the southeastern brackish water populations (NAK-B, KAR-B and ISH-B) had an effective population size (Ne) of 4,100. During the subsequent Yolandia Sea period (phase II), when the Baltic Sea had low salinity brackish water, Ne decreased to around 1,000, likely reflecting a founder effect during the colonisation of the newly available habitat. Throughout the Ancylus Lake (freshwater), the Initial Littorina Sea (increasing salinity) and the Littorina Sea (high salinity brackish water) periods (phases III-VI) Ne remained low around 1,000, probably due to competition from freshwater adapted populations during the Ancylus period and limited habitat in the Littorina period. The exact salinity of the Littorina Sea has been debated, but most studies indicate that it was 25-75% higher than at present (Weckström et al., 2017). In the western Baltic Sea, brackish water perch are currently living close to their maximum salinity tolerance of around 18 ppt (Christensen et al., 2019), hence the higher salinity of the Littorina Sea likely restricted perch to areas close to river mouths where salinities were locally lower. Zooarchaeological findings suggest that perch did persist in the western Baltic Sea during the Littorina Sea period (Schmölcke & Ritchie, 2010), but our demographic simulation suggests that this was in smaller and or more isolated populations than at present. Finally, as environmental conditions of the Baltic Sea changed from high salinity brackish water to the contemporary brackish/freshwater salinity levels during the transition from the Littorina Sea to the Post-Littorina Sea about 4.5-3 kya (Emeis et al., 2003), the western Baltic Sea perch saw a substantial increase in Ne to 4,400, possibly reflecting an increase in population connectivity and available habitat similar to the current Baltic Sea conditions.

### Implications for perch management

Our results are of special interest for management of perch, especially given that brackish water populations in the Baltic Sea are currently under pressure from several abiotic and biotic factors, including human perturbations of the environment. During the autumn, when western Baltic brackish water perch are returning to their natal streams, influxes of high-salinity sea water exceeding their physiological tolerance can lead to mass mortality among perch (Berg, 2012). There is also an increasing threat from cormorant predation during the winter, when the brackish water perch are congregating (Salmi et al., 2014; Veneranta et al., 2020) and in some areas of the Baltic Sea there is egg predation from three spined sticklebacks (Donadi et al., 2020). These pressures, combined with unregulated commercial and leisure harvesting during the summer, when the brackish water perch are in the marine environment, has led to greatly fluctuating population sizes of brackish water perch (Lindvig & Ebert, 2012). In addition, many streams in the western Baltic Sea have floodgates installed to hinder flooding of houses, which may deter the migration of perch and other migratory species of fish; in particular under future scenarios of climate change and rising sea levels. Particularly vulnerable populations may include ISK-B of only a few thousand individuals (Christensen et al., 2021), however for most populations there is no information of population sizes, which makes it difficult to identify at-risk populations. The genomic adaptations of southeastern brackish water populations indicate that they may not be easily rescued by migrants from upstream freshwater localities, but would have to await immigration from other brackish water localities. In isolated populations such as RAN-B, ROS-B and KET-B, local extinction would likely result in permanent disappearance of the brackish water ecotype making local nature management even more pertinent.

## Acknowledgement

The authors would like to thank Lasse Vinner, Pernille S. Olsen and Tina B. Brand at Globe Institute for help during laboratory work and professor Kim Præbel for valuable input. E.A.F.C. was supported by the Carlsberg Foundation (grant number CF19-0400). M.T.O. was supported via the BONUS BALTHEALTH project, BONUS (Art. 185), funded jointly by the EU, Innovation Fund Denmark (grants 6180-00001B and 6180-00002B), Forschungszentrum Jülich GmbH, German Federal Ministry of Education and Research (grant FKZ 03F0767A), Academy of Finland (grant 311966) and Swedish Foundation for Strategic Environmental Research (MISTRA). P.R.M. and H.C. were supported by Aage V. Jensens nature Foundation (Grant number 100307).

## Author contribution

MS, MTO and PRM conceived the study. MTPG, MTO and PRM provided funding. MS, MAK, PRM and HC collected the samples. MS, GP, EFC, THH and FGV performed the analyses. MS, GP, EFC and MTO drafted the manuscript with input from the remaining authors. All authors approved of the final version.

## Data availability

All resequencing data is publicly available at the European Nucleotide Archive (ENA), and can be accessed through the Project Number: **Will be added upon acceptance**.

All codes required to reproduce the computational analyses are available from GitHub (https://github.com/g-pacheco/PerchGenomics).

## Supplementary figures legends

Figure S1. Heatmap of depth of coverage for the 190 perch samples and the two blanks. Samples excluded from further analyses are labelled “BadSamples” and indicated by a purple square in the top row while retained samples are labelled “GoodSamples” and indicated by a green square.

Figure S2. Global Depth distribution. Density plot of the Global Depth calculated across all 190 perch samples. The purple vertical dashed line indicates the cutoff used, which was a maximum of 190 times the number of individuals.

## References

Ashburner, M., Ball, C. A., Blake, J. A., Botstein, D., Butler, H., Cherry, J. M., Davis, A. P., Dolinski, K., Dwight, S. S., Eppig, J. T., Harris, M. A., Hill, D. P., Issel-Tarver, L., Kasarskis, A., Lewis, S., Matese, J. C., Richardson, J. E., Ringwald, M., Rubin, G. M., & Sherlock, G. (2000). Gene Ontology: tool for the unification of biology. Nature Genetics, 25(1), 25–29.

Becht, E., McInnes, L., Healy, J., Dutertre, C.-A., Kwok, I. W. H., Ng, L. G., Ginhoux, F., & Newell, E. W. (2018). Dimensionality reduction for visualizing single-cell data using UMAP. Nature Biotechnology, 1(37), 38–45.

Behrmann-Godel, J., & Gerlach, G. (2008). First evidence for postzygotic reproductive isolation between two populations of Eurasian perch (Perca fluviatilis L.) within Lake Constance. Frontiers in Zoology, 5(3). https://doi.org/10.1186/1742-9994-5-3

Behrmann-Godel, J., Gerlach, G., & Eckmann, R. (2006). Kin and population recognition in sympatric Lake Constance perch (Perca fluviatilis L.): can assortative shoaling drive population divergence? Behavioral Ecology and Sociobiology, 59(4), 461–468.

Bekkevold, D., Jacobsen, L., Hemmer-Hansen, J., Berg, S., & Skov, C. (2015). From regionally predictable to locally complex population structure in a freshwater top predator: river systems are not always the unit of connectivity in Northern Pike Esox lucius. Ecology of Freshwater Fish, 24(2), 305–316.

Bergek, S., & Björklund, M. (2007). Cryptic barriers to dispersal within a lake allow genetic differentiation of Eurasian perch. Evolution; International Journal of Organic Evolution, 61(8), 2035–2041.

Bergek, S., & Björklund, M. (2009). Genetic and morphological divergence reveals local subdivision of perch (Perca fluviatilis L.). Biological Journal of the Linnean Society. Linnean Society of London, 96, 746–758.

Berg, S. (2012). Aborre: Perca fluviatisis Linnaeus, 1758. In H. Carl & P. R. Møller (Eds.), Atlas Over Danske Ferskvandsfisk. Staten naturhistoriske Museum, Københavns Universitet.

Björck, S. (1995). A review of the history of the Baltic Sea, 13.0-8.0 ka BP. Quaternary International: The Journal of the International Union for Quaternary Research, 27, 19–40.

Brauner, C. J., Shrimpton, J. M., & Randall, J. D. (1992). Effect of Short-Duration Seawater Exposure on Plasma Ion Concentrations and Swimming Performance in Coho Salmon (Oncorhynchus kisutch) Parr. Canadian Journal of Fisheries and Aquatic Sciences, 49, 2399–24635.

Brett, J. R. (1979). Physiological energetics. In Hoar, W. S. & Randall, D. J. (Ed.), Fish Physiology, Vol. VIII (pp. 279–352). Academic Press, New York, NY.

Carbon, S., Ireland, A., Mungall, C. J., Shu, S., Marshall, B., Lewis, S., AmiGO Hub, & Web Presence Working Group. (2009). AmiGO: online access to ontology and annotation data. Bioinformatics, 25(2), 288–289.

Christensen, E. A. F., Grosell, M., & Steffensen, J. F. (2019). Maximum salinity tolerance and osmoregulatory capabilities of European perch Perca fluviatilis populations originating from different salinity habitats. Conservation Physiology, 7(1), coz004.

Christensen, E. A. F., Skovrind, M., Olsen, M. T., Carl, H., Gravlund, P., & Møller, P. R. (2016). Hatching success in brackish water of Perca fluviatilis eggs obtained from the western Baltic Sea. Cybium, 6, 133–138.

Christensen, E. A. F., Svendsen, M. B. S., & Steffensen, J. F. (2017). Plasma osmolality and oxygen consumption of perch Perca fluviatilis in response to different salinities and temperatures. Journal of Fish Biology, 90(3), 819–833.

Christensen, E. A. F., Svendsen, M. B. S., & Steffensen, J. F. (2021). Population ecology, growth, and physico-chemical habitat of anadromous European perch Perca fluviatilis. Estuarine, Coastal and Shelf Science, 249, 107091.

Cock, P. J. A., Antao, T., Chang, J. T., Chapman, B. A., Cox, C. J., Dalke, A., Friedberg, I., Hamelryck, T., Kauff, F., Wilczynski, B., & de Hoon, M. J. L. (2009). Biopython: freely available Python tools for computational molecular biology and bioinformatics. Bioinformatics, 25(11), 1422–1423.

Cole, D. M. (2018). Responses to lowered salinity in the Pacific spiny dogfish, Squalus suckleyi, a marginally euryhaline shark (G. Goss (ed.)) [Master of Science In Physiology, Cell and Developmental Biology]. University of Alberta.

Cui, J., Liu, J., Li, J., Cheng, D., & Dai, C. (2022). Genome-wide sequence identification and expression analysis of N6 -methyladenosine demethylase in sugar beet (Beta vulgaris L.) under salt stress. PeerJ, 10, e12719.

Culling, M. A., Janko, K., Boron, A., Vasil’ev, V. P., Côté, I. M., & Hewitt, G. M. (2006). European colonization by the spined loach (Cobitis taenia) from Ponto-Caspian refugia based on mitochondrial DNA variation. Molecular Ecology, 15(1), 173–190.

Deane, E. E., & Woo, N. Y. S. (2011). Advances and perspectives on the regulation and expression of piscine heat shock proteins. Reviews in Fish Biology and Fisheries, 21(2), 153–185.

Donadi, S., Bergström, L., Bertil Berglund, J. M., Anette, B., Mikkola, R., Saarinen, A., & Bergström, U. (2020). Perch and pike recruitment in coastal bays limited by stickleback predation and environmental forcing. Estuarine, Coastal and Shelf Science, 246, 107052.

El-Shewy, H. M., & Luttrell, L. M. (2009). Chapter 24 Insulin-Like Growth Factor-2/Mannose-6 Phosphate Receptors. In Vitamins & Hormones (Vol. 80, pp. 667–697). Academic Press.

Elshire, R. J., Glaubitz, J. C., Sun, Q., Poland, J. A., Kawamoto, K., Buckler, E. S., & Mitchell, S. E. (2011). A robust, simple genotyping-by-sequencing (GBS) approach for high diversity species. PloS One, 6(5), e19379.

Emeis, K.-C., Struck, U., Blanz, T., Kohly, A., & Voβ, M. (2003). Salinity changes in the central Baltic Sea (NW Europe) over the last 10000 years. Holocene, 13(3), 411–421.

Evans, D. H. (1984). The Roles of Gill Permeability and Transport Mechanisms in Euryhalinity. In Fish Physiology (pp. 239–283). cademic Press.

Feder, J. L., Egan, S. P., & Nosil, P. (2012). The genomics of speciation-with-gene-flow. Trends in Genetics: TIG, 28(7), 342–350.

Foote, A. D., Vijay, N., Ávila-Arcos, M. C., Baird, R. W., Durban, J. W., Fumagalli, M., Gibbs, R. A., Hanson, M. B., Korneliussen, T. S., Martin, M. D., Robertson, K. M., Sousa, V. C., Vieira, F. G., Vinař, T., Wade, P., Worley, K. C., Excoffier, L., Morin, P. A., Gilbert, M. T. P., & Wolf, J. B. W. (2016). Genome-culture coevolution promotes rapid divergence of killer whale ecotypes. Nature Communications, 7, 11693.

Froese, R., & Binohlan, C. (2000). Empirical relationships to estimate asymptotic length, length at first maturity and length at maximum yield per recruit in fishes, with a simple method to evaluate length frequency data. Journal of Fish Biology, 56(4), 758–773.

Hall, A., & van Boeckel, M. (2020). Origin of the Baltic Sea basin by Pleistocene glacial erosion. GFF, 142(3), 237–252.

Hansen, C. C. R., Hvilsom, C., Schmidt, N. M., Aastrup, P., Van Coeverden de Groot, P. J., Siegismund, H. R., & Heller, R. (2018). The Muskox Lost a Substantial Part of Its Genetic Diversity on Its Long Road to Greenland. Current Biology: CB, 28(24), 4022–4028.e5.

Hohenlohe, P. A., Bassham, S., Etter, P. D., Stiffler, N., Johnson, E. A., & Cresko, W. A. (2010). Population genomics of parallel adaptation in threespine stickleback using sequenced RAD tags. PLoS Genetics, 6(2), e1000862.

Jacobsen, L., Bekkevold, D., Berg, S., Jepsen, N., Koed, A., Aarestrup, K., Baktoft, H., & Skov, C. (2017). Pike (Esox lucius L.) on the edge: consistent individual movement patterns in transitional waters of the western Baltic. Hydrobiologia, 784(1), 143–154.

Johnson, M., Zaretskaya, I., Raytselis, Y., Merezhuk, Y., McGinnis, S., & Madden, T. L. (2008). NCBI BLAST: a better web interface. Nucleic Acids Research, 36(Web Server issue), W5–W9.

Kolde, R. (2012). Pheatmap: pretty heatmaps (R package) [Computer software]. https://github.com/raivokolde/pheatmap

Kontula, T., & Väinölä, R. (2001). Postglacial colonization of Northern Europe by distinct phylogeographic lineages of the bullhead, Cottus gobio. Molecular Ecology, 10(8), 1983–2002.

Korneliussen, T. S., Albrechtsen, A., & Nielsen, R. (2014). ANGSD: Analysis of Next Generation Sequencing Data. BMC Bioinformatics, 15, 356.

Langerhans, R. B., Brian Langerhans, R., & Riesch, R. (2013). Speciation by selection: A framework for understanding ecology’s role in speciation. Current Zoology, 59(1), 31–52.

Larsen, E. H., Deaton, L. E., Onken, H., O’Donnell, M., Grosell, M., Dantzler, W. H., & Weihrauch, D. (2014). Osmoregulation and excretion. Comprehensive Physiology, 4(2), 405–573.

Larsen, P. F., Nielsen, E. E., Williams, T. D., & Loeschcke, V. (2008). Intraspecific variation in expression of candidate genes for osmoregulation, heme biosynthesis and stress resistance suggests local adaptation in European flounder (Platichthys flesus). Heredity, 101(3), 247–259.

Leppäranta, M., & Myrberg, K. (2009). Physical Oceanography of the Baltic Sea. Springer Science & Business Media.

Li, H., Handsaker, B., Wysoker, A., Fennell, T., Ruan, J., Homer, N., Marth, G., Abecasis, G., Durbin, R., & 1000 Genome Project Data Processing Subgroup. (2009). The Sequence Alignment/Map format and SAMtools. Bioinformatics, 25(16), 2078–2079.

Lindvig, D., & Ebert, K. M. (2012). Tilstanden og udviklingspotentialet hos brakvandsgedder og -aborrer i farvandet omkring Sydsjælland, Møn og Lolland-Falster. Danmarks Sportsfiskerforbund.

Li, N., Huang, Z., Ding, L., Shi, H., & Hong, M. (2021). Endoplasmic reticulum unfolded protein response modulates the adaptation of Trachemys scripta elegans in salinity water. Comparative Biochemistry and Physiology. Toxicology & Pharmacology: CBP, 248, 109102.

Li, S., He, F., Wen, H., Si, Y., Liu, M., Huang, Y., & Wu, S. (2020). Half Smooth Tongue Sole (Cynoglossus semilaevis) Under Low Salinity Stress Can Change Hepatic igf2 Expression Through DNA Methylation. Journal of Ocean University of China, 19(1), 171–182.

Liu, S., Hansen, M. M., & Jacobsen, M. W. (2016). Region-wide and ecotype-specific differences in demographic histories of threespine stickleback populations, estimated from whole genome sequences. Molecular Ecology, 25(20), 5187–5202.

Liu, X., & Fu, Y.-X. (2020). Stairway Plot 2: demographic history inference with folded SNP frequency spectra. Genome Biology, 21(1), 280.

Lutz, P. L. (1972). Ionic and body compartment responses to increasing salinity in the perch Perca fluviatilis. Comparative Biochemistry and Physiology. A, Comparative Physiology, 42(3), 711–717.

Magurran, A. E., Khachonpisitsak, S., & Ahmad, A. B. (2011). Biological diversity of fish communities: pattern and process. Journal of Fish Biology, 79, 1393–1412.

Mas-Sandoval, A., Pope, N. S., Nielsen, K. N., Altinkaya, I., Fumagalli, M., & Korneliussen, T. S. (2022). Fast and accurate estimation of multidimensional site frequency spectra from low-coverage high-throughput sequencing data. GigaScience, 11, 1–9.

McDonald, J. F., & Ayala, F. J. (1974). Genetic response to environmental heterogeneity. Nature, 250(467), 572–574.

Meisner, J., & Albrechtsen, A. (2018). Inferring Population Structure and Admixture Proportions in Low-Depth NGS Data. Genetics, 210(2), 719–731.

Nei, M., & Tajima, F. (1981). Genetic drift and estimation of effective population size. Genetics, 98(3), 625–640.

Nesbø, C. L., Fossheim, T., Vollestad, L. A., & Jakobsen, K. S. (1999). Genetic divergence and phylogeographic relationships among European perch (Perca fluviatilis) populations reflect glacial refugia and postglacial colonization. Molecular Ecology, 8(9), 1387–1404.

Norman, J. D., Danzmann, R. G., Glebe, B., & Ferguson, M. M. (2011). The genetic basis of salinity tolerance traits in Arctic charr (Salvelinus alpinus). BMC Genetics, 12(1), 81.

Olsson, J., Mo, K., Florin, A.-B., Aho, T., & Ryman, N. (2011). Genetic population structure of perch Perca fluviatilis along the Swedish coast of the Baltic Sea. Journal of Fish Biology, 79(1), 122–137.

Özmen, S., & Tabur, S. (2020). Functions of folic acid (Vitamin B9) against cytotoxic effects of salt stress in Hordeum vulgare L. Pakistan Journal of Botany, 52(1).

Radersma, R., Noble, D. W. A., & Uller, T. (2020). Plasticity leaves a phenotypic signature during local adaptation. Evolution Letters, 4(4), 360–370.

Remane, A. (1934). Die Gastrotrichen des Küstengrundwassers von Schilksee. Aus Den Schriften Des Naturwissenschaftlichen Vereins Für Schleswig-Holstein, 20, 473–478.

Salmi, J. A., Auvinen, H., Raitaniemi, J., Kurkilahti, M., Lilja, J., & Maikola, R. (2014). Perch (Perca fluviatilis) and pikeperch (Sander lucioperca) in the diet of the great cormorant (Phalacrocorax carbo) and effects on catches in the Archipelago Sea, Southwest coast of Finland. Fisheries Research, 164, 26–34.

Schmölcke, U., & Ritchie, K. (2010). A new method in palaeoecology: fish community structure indicates environmental changes. International Journal of Earth Sciences, 99(8), 1763–1772.

Seehausen, O., Butlin, R. K., Keller, I., Wagner, C. E., Boughman, J. W., Hohenlohe, P. A., Peichel, C. L., Saetre, G.-P., Bank, C., Brännström, A., Brelsford, A., Clarkson, C. S., Eroukhmanoff, F., Feder, J. L., Fischer, M. C., Foote, A. D., Franchini, P., Jiggins, C. D., Jones, F. C., … Widmer, A. (2014). Genomics and the origin of species. Nature Reviews. Genetics, 15(3), 176–192.

Skotte, L., Korneliussen, T. S., & Albrechtsen, A. (2013). Estimating individual admixture proportions from next generation sequencing data. Genetics, 195(3), 693–702.

Skovrind, M., Christensen, E. A. F., Carl, H., Jacobsen, L., & Møller, P. R. (2013). Marine spawning sites of perch Perca fluviatilis revealed by oviduct-inserted acoustic transmitters. Aquatic Biology, 19, 201–206.

Skovrind, M., Olsen, M. T., Vieira, F. G., Pacheco, G., Carl, H., Gilbert, M. T. P., & Møller, P. R. (2016). Genomic population structure of freshwater-resident and anadromous ide (Leuciscus idus) in north-western Europe. Ecology and Evolution, 6(4), 1064–1074.

Sørensen, J. G., Kristensen, T. N., & Loeschcke, V. (2003). The evolutionary and ecological role of heat shock proteins. Ecology Letters, 6(11), 1025–1037.

Sunde, J., Yıldırım, Y., Tibblin, P., Bekkevold, D., Skov, C., Nordahl, O., Larsson, P., & Forsman, A. (2022). Drivers of neutral and adaptive differentiation in pike (Esox lucius) populations from contrasting environments. Molecular Ecology, 31(4), 1093–1110.

The Gene Ontology Consortium. (2021). The Gene Ontology resource: enriching a GOld mine. Nucleic Acids Research, 49, 325–334.

Toomey, L., Dellicour, S., Vanina, T., Pegg, J., Kaczkowski, Z., Kouřil, J., Teletchea, F., Bláha, M., Fontaine, P., & Lecocq, T. (2020). Getting off on the right foot: Integration of spatial distribution of genetic variability for aquaculture development and regulations, the European perch case. Aquaculture, 521, 734981.

Veneranta, L., Heikinheimo, O., & Marjomäki, T. J. (2020). Cormorant (Phalacrocorax carbo) predation on a coastal perch (Perca fluviatilis) population: estimated effects based on PIT tag mark-recapture experiment. ICES Journal of Marine Science, 77(7-8), 2611–2622.

Wang, W., Li, W., Cheng, Z., Sun, J., Gao, J., Li, J., Niu, X., Amjid, M. W., Yang, H., Zhu, G., Zhang, D., & Guo, W. (2022). Transcriptome-wide N6-methyladenosine profiling of cotton root provides insights for salt stress tolerance. Environmental and Experimental Botany, 194, 104729.

Ward, R. D., Woodwark, M., & Skibinski, D. O. F. (1994). A comparison of genetic diversity levels in marine, freshwater, and anadromous fishes. Journal of Fish Biology, 44(2), 213–232.

Weckström, K., Lewis, J. P., Andrén, E., Ellegaard, M., Rasmussen, P., Ryves, D. B., & Telford, R. (2017). Palaeoenvironmental History of the Baltic Sea: One of the Largest Brackish-Water Ecosystems in the World. In K. Weckström, K. M. Saunders, P. A. Gell, & C. G. Skilbeck (Eds.), Applications of Paleoenvironmental Techniques in Estuarine Studies (pp. 615–662). Springer Netherlands.

Whitfield, A. K., Elliott, M., Basset, A., Blaber, S. J. M., & West, R. J. (2012). Paradigms in estuarine ecology – A review of the Remane diagram with a suggested revised model for estuaries. Estuarine, Coastal and Shelf Science, 97, 78–90.

Wickham, H. (2016). ggplot2: Elegant Graphics for Data Analysis. Springer.

Wright, S. (1943). Isolation by Distance. Genetics, 28(2), 114–138.

Zheng, H., Sun, X., Li, J., Song, Y., Song, J., Wang, F., Liu, L., Zhang, X., & Sui, N. (2021). Analysis of N6-methyladenosine reveals a new important mechanism regulating the salt tolerance of sweet sorghum. Plant Science: An International Journal of Experimental Plant Biology, 304, 110801.

