## Supplementary material for "Genomic differentiation among European perch in the western Baltic Sea reflects colonisation history and local adaptation": Table 1

Table1. European perch sample localities and their abbreviations.

| Abbreviation | Locality name | Sample size | Environment |
|---|---|---|---|
| TAN-F | Tange Sø | 19 | Fresh water |
| RAN-B | Randers Fjord | 30 | Brackish |
| FAR-F | Fårup Sø | 10 | Fresh water |
| SJA-F | Sjælsø | 12 | Fresh water |
| SON-F | Sønder Sø | 13 | Fresh water |
| TYB-F | Tystrup-Bavelse Sø | 13 | Fresh water |
| POL-F | Pøle Å | 18 | Fresh water |
| ROS-B | Roskilde Fjord | 12 | Brackish |
| KET-B | kettinge Nor | 13 | Brackish |
| NAK-B | Nakskov Inderfjord | 12 | Brackish |
| KAR-B | Karrebæk Fjord | 20 | Brackish |
| ISH-B | Ishøj Havn | 18 | Brackish |
