## Supplementary material for "Genomic differentiation among European perch in the western Baltic Sea reflects colonisation history and local adaptation": Table S1

| SORT | Sample ID | Sampling date | Species | Locality | Locality-abbrev. | Lab_Nr. | Zmuc_Nr. | Extraction_date | Life_stage | Environment | Country | Sample_type | Plate_ID | Flowcell | Lane | Barcode | fastq_file |
|---|---|---|---|---|---|---|---|---|---|---|---|---|---|---|---|---|---|
| 1 | ROS-B_01 | 24/09 20013 | Perca_fluviatilis | Roskilde Fjord | ROS-B | 1 | P45545 | 1/21/2014 | Adult | Brackish | Denmark | Fin clipping | PGP_1 | X | 5 | TTGA | PGP-C57G8ANXX_5_fastq.gz |
| 2 | ROS-B_02 | 24/09 20013 | Perca_fluviatilis | Roskilde Fjord | ROS-B | 2 | P45546 | 1/21/2014 | Adult | Brackish | Denmark | Fin clipping | PGP_1 | X | 5 | CTAGAA | PGP-C57G8ANXX_5_fastq.gz |
| 3 | ROS-B_03 | 24/09 20013 | Perca_fluviatilis | Roskilde Fjord | ROS-B | 3 | P45547 | 1/21/2014 | Adult | Brackish | Denmark | Fin clipping | PGP_1 | X | 5 | CGTAGGA | PGP-C57G8ANXX_5_fastq.gz |
| 4 | ROS-B_04 | 24/09 20013 | Perca_fluviatilis | Roskilde Fjord | ROS-B | 4 | P45548 | 2/25/2014 | Adult | Brackish | Denmark | Fin clipping | PGP_1 | X | 5 | TGGTCAG | PGP-C57G8ANXX_5_fastq.gz |
| 5 | ROS-B_05 | 24/09 20013 | Perca_fluviatilis | Roskilde Fjord | ROS-B | 5 | P45549 | 2/25/2014 | Adult | Brackish | Denmark | Fin clipping | PGP_1 | X | 5 | TTAACGA | PGP-C57G8ANXX_5_fastq.gz |
| 6 | ROS-B_06 | 24/09 20013 | Perca_fluviatilis | Roskilde Fjord | ROS-B | 6 | P45550 | 2/25/2014 | Adult | Brackish | Denmark | Fin clipping | PGP_1 | X | 5 | ACTCGACA | PGP-C57G8ANXX_5_fastq.gz |
| 7 | ROS-B_07 | 24/09 20013 | Perca_fluviatilis | Roskilde Fjord | ROS-B | 7 | P45551 | 2/25/2014 | Adult | Brackish | Denmark | Fin clipping | PGP_1 | X | 5 | GCCATACG | PGP-C57G8ANXX_5_fastq.gz |
| 8 | ROS-B_08 | 24/09 20013 | Perca_fluviatilis | Roskilde Fjord | ROS-B | 8 | P45552 | 2/25/2014 | Adult | Brackish | Denmark | Fin clipping | PGP_1 | X | 5 | CTCCGACG | PGP-C57G8ANXX_5_fastq.gz |
| 9 | ROS-B_09 | 24/09 20013 | Perca_fluviatilis | Roskilde Fjord | ROS-B | 9 | P45553 | 2/25/2014 | Adult | Brackish | Denmark | Fin clipping | PGP_1 | X | 5 | GGTA | PGP-C57G8ANXX_5_fastq.gz |
| 10 | ROS-B_10 | 24/09 20013 | Perca_fluviatilis | Roskilde Fjord | ROS-B | 10 | P45554 | 2/25/2014 | Adult | Brackish | Denmark | Fin clipping | PGP_1 | X | 5 | GATACG | PGP-C57G8ANXX_5_fastq.gz |
| 11 | ROS-B_11 | 24/09 20013 | Perca_fluviatilis | Roskilde Fjord | ROS-B | 11 | P45555 | 2/25/2014 | Adult | Brackish | Denmark | Fin clipping | PGP_1 | X | 5 | GATAGCG | PGP-C57G8ANXX_5_fastq.gz |
| 12 | ROS-B_12 | 24/09 20013 | Perca_fluviatilis | Roskilde Fjord | ROS-B | 12 | P45556 | 2/25/2014 | Adult | Brackish | Denmark | Fin clipping | PGP_1 | X | 5 | GACGTGA | PGP-C57G8ANXX_5_fastq.gz |
| 13 | KAR-B_01 | 02/10 2013 | Perca_fluviatilis | Karrebæk Fjord | KAR-B | 13 | P45581 | 41731 | Adult | Brackish | Denmark | Fin clipping | PGP_1 | X | 5 | TAGCGCG | PGP-C57G8ANXX_5_fastq.gz |
| 14 | KAR-B_02 | 02/10 2013 | Perca_fluviatilis | Karrebæk Fjord | KAR-B | 14 | P45582 | 41731 | Adult | Brackish | Denmark | Fin clipping | PGP_1 | X | 5 | TGAGAGCA | PGP-C57G8ANXX_5_fastq.gz |
| 15 | KAR-B_03 | 02/10 2013 | Perca_fluviatilis | Karrebæk Fjord | KAR-B | 15 | P45583 | 41731 | Adult | Brackish | Denmark | Fin clipping | PGP_1 | X | 5 | ATTCAGCA | PGP-C57G8ANXX_5_fastq.gz |
| 16 | KAR-B_04 | 02/10 2013 | Perca_fluviatilis | Karrebæk Fjord | KAR-B | 16 | P45584 | 41731 | Adult | Brackish | Denmark | Fin clipping | PGP_1 | X | 5 | TCTGATAA | PGP-C57G8ANXX_5_fastq.gz |
| 17 | KAR-B_05 | 02/10 2013 | Perca_fluviatilis | Karrebæk Fjord | KAR-B | 17 | P45585 | 41731 | Adult | Brackish | Denmark | Fin clipping | PGP_1 | X | 5 | TCTGG | PGP-C57G8ANXX_5_fastq.gz |
| 18 | KAR-B_06 | 02/10 2013 | Perca_fluviatilis | Karrebæk Fjord | KAR-B | 18 | P45586 | 41731 | Adult | Brackish | Denmark | Fin clipping | PGP_1 | X | 5 | TCCGAA | PGP-C57G8ANXX_5_fastq.gz |
| 19 | KAR-B_07 | 02/10 2013 | Perca_fluviatilis | Karrebæk Fjord | KAR-B | 19 | P45587 | 41731 | Adult | Brackish | Denmark | Fin clipping | PGP_1 | X | 5 | ACCTCGA | PGP-C57G8ANXX_5_fastq.gz |
| 20 | KAR-B_08 | 02/10 2013 | Perca_fluviatilis | Karrebæk Fjord | KAR-B | 20 | P45588 | 41731 | Adult | Brackish | Denmark | Fin clipping | PGP_1 | X | 5 | CTAAGAA | PGP-C57G8ANXX_5_fastq.gz |
| 21 | KAR-B_09 | 02/10 2013 | Perca_fluviatilis | Karrebæk Fjord | KAR-B | 21 | P45589 | 41731 | Adult | Brackish | Denmark | Fin clipping | PGP_1 | X | 5 | GCGTTAA | PGP-C57G8ANXX_5_fastq.gz |
| 22 | KAR-B_10 | 02/10 2013 | Perca_fluviatilis | Karrebæk Fjord | KAR-B | 22 | P45590 | 41731 | Adult | Brackish | Denmark | Fin clipping | PGP_1 | X | 5 | GTGTGCCA | PGP-C57G8ANXX_5_fastq.gz |
| 23 | KAR-B_11 | 02/10 2013 | Perca_fluviatilis | Karrebæk Fjord | KAR-B | 23 | P45591 | 2/13/2014 | Adult | Brackish | Denmark | Fin clipping | PGP_1 | X | 5 | TAGTGCCG | PGP-C57G8ANXX_5_fastq.gz |
| 24 | KAR-B_12 | 02/10 2013 | Perca_fluviatilis | Karrebæk Fjord | KAR-B | 24 | P45592 | 2/13/2014 | Adult | Brackish | Denmark | Fin clipping | PGP_1 | X | 5 | AGGATCCG | PGP-C57G8ANXX_5_fastq.gz |
| 25 | PGP_1-Blank | PGP_1-Blank | Na | Na | Na | Na | Na | Na | Na | Na | Na | Na | PGP_1 | X | 5 | CGGCA | PGP-C57G8ANXX_5_fastq.gz |
| 26 | RAN-B_01 | 08/10 2013 | Perca_fluviatilis | Randers Fjord | RAN-B | 25 | P45601 | 2/13/2014 | Adult | Brackish | Denmark | Fin clipping | PGP_1 | X | 5 | CAGTGA | PGP-C57G8ANXX_5_fastq.gz |
| 27 | RAN-B_02 | 08/10 2013 | Perca_fluviatilis | Randers Fjord | RAN-B | 26 | P45603 | 2/13/2014 | Adult | Brackish | Denmark | Fin clipping | PGP_1 | X | 5 | TTGCTAA | PGP-C57G8ANXX_5_fastq.gz |
| 28 | RAN-B_03 | 08/10 2013 | Perca_fluviatilis | Randers Fjord | RAN-B | 27 | P45604 | 2/13/2014 | Adult | Brackish | Denmark | Fin clipping | PGP_1 | X | 5 | GGTCACG | PGP-C57G8ANXX_5_fastq.gz |
| 29 | RAN-B_04 | 08/10 2013 | Perca_fluviatilis | Randers Fjord | RAN-B | 28 | P45605 | 41695 | Adult | Brackish | Denmark | Fin clipping | PGP_1 | X | 5 | CTCAAGA | PGP-C57G8ANXX_5_fastq.gz |
| 30 | RAN-B_05 | 08/10 2013 | Perca_fluviatilis | Randers Fjord | RAN-B | 29 | P45606 | 41695 | Adult | Brackish | Denmark | Fin clipping | PGP_1 | X | 5 | AAGACTCA | PGP-C57G8ANXX_5_fastq.gz |
| 31 | RAN-B_06 | 08/10 2013 | Perca_fluviatilis | Randers Fjord | RAN-B | 30 | P45607 | 41695 | Adult | Brackish | Denmark | Fin clipping | PGP_1 | X | 5 | GGCTATCA | PGP-C57G8ANXX_5_fastq.gz |
| 32 | RAN-B_07 | 08/10 2013 | Perca_fluviatilis | Randers Fjord | RAN-B | 31 | P45608 | 41695 | Adult | Brackish | Denmark | Fin clipping | PGP_1 | X | 5 | GAATACAA | PGP-C57G8ANXX_5_fastq.gz |
| 33 | RAN-B_08 | 08/10 2013 | Perca_fluviatilis | Randers Fjord | RAN-B | 32 | P45609 | 41695 | Adult | Brackish | Denmark | Fin clipping | PGP_1 | X | 5 | GAAGA | PGP-C57G8ANXX_5_fastq.gz |
| 34 | RAN-B_09 | 08/10 2013 | Perca_fluviatilis | Randers Fjord | RAN-B | 33 | P45610 | 41695 | Adult | Brackish | Denmark | Fin clipping | PGP_1 | X | 5 | AGTCCG | PGP-C57G8ANXX_5_fastq.gz |
| 35 | RAN-B_10 | 08/10 2013 | Perca_fluviatilis | Randers Fjord | RAN-B | 34 | P45612 | 41695 | Adult | Brackish | Denmark | Fin clipping | PGP_1 | X | 5 | TGAGGCG | PGP-C57G8ANXX_5_fastq.gz |
| 36 | RAN-B_11 | 08/10 2013 | Perca_fluviatilis | Randers Fjord | RAN-B | 35 | P45613 | 41695 | Adult | Brackish | Denmark | Fin clipping | PGP_1 | X | 5 | TCGTAGA | PGP-C57G8ANXX_5_fastq.gz |
| 37 | RAN-B_12 | 08/10 2013 | Perca_fluviatilis | Randers Fjord | RAN-B | 36 | P45614 | 41695 | Adult | Brackish | Denmark | Fin clipping | PGP_1 | X | 5 | CGTGGAG | PGP-C57G8ANXX_5_fastq.gz |
| 38 | NAK-B_01 | 27/10 2013 | Perca_fluviatilis | Nakskov indrefjord | NAK-B | 37 | p45561 | 2/13/2014 | Adult | Brackish | Denmark | Fin clipping | PGP_1 | X | 5 | CCACTACG | PGP-C57G8ANXX_5_fastq.gz |
| 39 | NAK-B_02 | 27/10 2013 | Perca_fluviatilis | Nakskov indrefjord | NAK-B | 38 | p45562 | 2/13/2014 | Adult | Brackish | Denmark | Fin clipping | PGP_1 | X | 5 | CCTGCACG | PGP-C57G8ANXX_5_fastq.gz |
| 40 | NAK-B_03 | 27/10 2013 | Perca_fluviatilis | Nakskov indrefjord | NAK-B | 39 | p45563 | 2/13/2014 | Adult | Brackish | Denmark | Fin clipping | PGP_1 | X | 5 | CTGACACG | PGP-C57G8ANXX_5_fastq.gz |
| 41 | NAK-B_04 | 27/10 2013 | Perca_fluviatilis | Nakskov indrefjord | NAK-B | 40 | p45564 | 41858 | Adult | Brackish | Denmark | Fin clipping | PGP_1 | X | 5 | ACGAG | PGP-C57G8ANXX_5_fastq.gz |
| 42 | NAK-B_05 | 27/10 2013 | Perca_fluviatilis | Nakskov indrefjord | NAK-B | 41 | p45565 | 41858 | Adult | Brackish | Denmark | Fin clipping | PGP_1 | X | 5 | GCATGG | PGP-C57G8ANXX_5_fastq.gz |
| 43 | NAK-B_06 | 27/10 2013 | Perca_fluviatilis | Nakskov indrefjord | NAK-B | 42 | p45566 | 41858 | Adult | Brackish | Denmark | Fin clipping | PGP_1 | X | 5 | GACTAGG | PGP-C57G8ANXX_5_fastq.gz |
| 44 | NAK-B_07 | 27/10 2013 | Perca_fluviatilis | Nakskov indrefjord | NAK-B | 43 | p45567 | 41858 | Adult | Brackish | Denmark | Fin clipping | PGP_1 | X | 5 | AGCTCCG | PGP-C57G8ANXX_5_fastq.gz |
| 45 | NAK-B_08 | 27/10 2013 | Perca_fluviatilis | Nakskov indrefjord | NAK-B | 44 | p45568 | 41858 | Adult | Brackish | Denmark | Fin clipping | PGP_1 | X | 5 | GTACGCG | PGP-C57G8ANXX_5_fastq.gz |
| 46 | NAK-B_09 | 27/10 2013 | Perca_fluviatilis | Nakskov indrefjord | NAK-B | 45 | p45569 | 41858 | Adult | Brackish | Denmark | Fin clipping | PGP_1 | X | 5 | GGTTCGCA | PGP-C57G8ANXX_5_fastq.gz |
| 47 | NAK-B_10 | 27/10 2013 | Perca_fluviatilis | Nakskov indrefjord | NAK-B | 46 | p45570 | 41858 | Adult | Brackish | Denmark | Fin clipping | PGP_1 | X | 5 | TTAAGCAA | PGP-C57G8ANXX_5_fastq.gz |
| 48 | NAK-B_11 | 27/10 2013 | Perca_fluviatilis | Nakskov indrefjord | NAK-B | 47 | p45571 | 41858 | Adult | Brackish | Denmark | Fin clipping | PGP_1 | X | 5 | TCCGTGAA | PGP-C57G8ANXX_5_fastq.gz |
| 49 | NAK-B_12 | 27/10 2013 | Perca_fluviatilis | Nakskov indrefjord | NAK-B | 48 | p45572 | 41858 | Adult | Brackish | Denmark | Fin clipping | PGP_1 | X | 5 | TTCTA | PGP-C57G8ANXX_5_fastq.gz |
| 50 | TAN-F_01 | 22/10 2013 | Perca_fluviatilis | Tange Sø | TAN-F | 49 | P45641 | 41731 | Juvenile | Fresh | Denmark | Fin clipping | PGP_1 | X | 5 | GTCCAA | PGP-C57G8ANXX_5_fastq.gz |
| 51 | TAN-F_02 | 22/10 2013 | Perca_fluviatilis | Tange Sø | TAN-F | 50 | P45642 | 41731 | Juvenile | Fresh | Denmark | Fin clipping | PGP_1 | X | 5 | ATAGCAA | PGP-C57G8ANXX_5_fastq.gz |
| 52 | TAN-F_03 | 22/10 2013 | Perca_fluviatilis | Tange Sø | TAN-F | 51 | P45643 | 41731 | Juvenile | Fresh | Denmark | Fin clipping | PGP_1 | X | 5 | TAACGAA | PGP-C57G8ANXX_5_fastq.gz |
| 53 | TAN-F_04 | 22/10 2013 | Perca_fluviatilis | Tange Sø | TAN-F | 52 | P45644 | 41731 | Juvenile | Fresh | Denmark | Fin clipping | PGP_1 | X | 5 | AACGCTA | PGP-C57G8ANXX_5_fastq.gz |
| 54 | TAN-F_05 | 22/10 2013 | Perca_fluviatilis | Tange Sø | TAN-F | 53 | P45645 | 41731 | Juvenile | Fresh | Denmark | Fin clipping | PGP_1 | X | 5 | ATCGACCG | PGP-C57G8ANXX_5_fastq.gz |
| 55 | TAN-F_06 | 22/10 2013 | Perca_fluviatilis | Tange Sø | TAN-F | 54 | P45646 | 41731 | Juvenile | Fresh | Denmark | Fin clipping | PGP_1 | X | 5 | GAGCTGCA | PGP-C57G8ANXX_5_fastq.gz |
| 56 | TAN-F_07 | 22/10 2013 | Perca_fluviatilis | Tange Sø | TAN-F | 55 | P45647 | 41731 | Juvenile | Fresh | Denmark | Fin clipping | PGP_1 | X | 5 | GATCGTCG | PGP-C57G8ANXX_5_fastq.gz |
| 57 | TAN-F_08 | 22/10 2013 | Perca_fluviatilis | Tange Sø | TAN-F | 56 | P45648 | 41731 | Juvenile | Fresh | Denmark | Fin clipping | PGP_1 | X | 5 | AGTCG | PGP-C57G8ANXX_5_fastq.gz |

| | | | | | | | | | | | | | | | | | |
|---|---|---|---|---|---|---|---|---|---|---|---|---|---|---|---|---|---|
| 58 | TAN-F_09 | 22/10 2013 | Perca_fluviatilis | Tange Sø | TAN-F | 57 | P45649 | 41731 | Juvenile | Fresh | Denmark | Fin clipping | PGP_1 | X | 5 | CAGGTG | PGP-C57G8ANXX_5_fastq.gz |
| 59 | TAN-F_10 | 22/10 2013 | Perca_fluviatilis | Tange Sø | TAN-F | 58 | P45650 | 41731 | Juvenile | Fresh | Denmark | Fin clipping | PGP_1 | X | 5 | CCTAACG | PGP-C57G8ANXX_5_fastq.gz |
| 60 | TAN-F_11 | 22/10 2013 | Perca_fluviatilis | Tange Sø | TAN-F | 59 | P45651 | 41731 | Juvenile | Fresh | Denmark | Fin clipping | PGP_1 | X | 5 | CTGACCG | PGP-C57G8ANXX_5_fastq.gz |
| 61 | TAN-F_12 | 22/10 2013 | Perca_fluviatilis | Tange Sø | TAN-F | 60 | P45652 | 41731 | Juvenile | Fresh | Denmark | Fin clipping | PGP_1 | X | 5 | TCTATGG | PGP-C57G8ANXX_5_fastq.gz |
| 62 | FAR-F_01 | 22/10 2013 | Perca_fluviatilis | Fårup Sø | FAR-F | 61 | P45621 | 2/13/2014 | Adult | Fresh | Denmark | Fin clipping | PGP_1 | X | 5 | TAACGACA | PGP-C57G8ANXX_5_fastq.gz |
| 63 | FAR-F_02 | 22/10 2013 | Perca_fluviatilis | Fårup Sø | FAR-F | 62 | P45622 | 2/13/2014 | Adult | Fresh | Denmark | Fin clipping | PGP_1 | X | 5 | AGACTAAG | PGP-C57G8ANXX_5_fastq.gz |
| 64 | FAR-F_03 | 22/10 2013 | Perca_fluviatilis | Fårup Sø | FAR-F | 63 | P45623 | 2/13/2014 | Adult | Fresh | Denmark | Fin clipping | PGP_1 | X | 5 | AGAGTCAA | PGP-C57G8ANXX_5_fastq.gz |
| 65 | FAR-F_04 | 22/10 2013 | Perca_fluviatilis | Fårup Sø | FAR-F | 64 | P45624 | 41858 | Adult | Fresh | Denmark | Fin clipping | PGP_1 | X | 5 | TACAGA | PGP-C57G8ANXX_5_fastq.gz |
| 66 | FAR-F_05 | 22/10 2013 | Perca_fluviatilis | Fårup Sø | FAR-F | 65 | P45625 | 41858 | Adult | Fresh | Denmark | Fin clipping | PGP_1 | X | 5 | TCACGA | PGP-C57G8ANXX_5_fastq.gz |
| 67 | FAR-F_06 | 22/10 2013 | Perca_fluviatilis | Fårup Sø | FAR-F | 66 | P45626 | 41858 | Adult | Fresh | Denmark | Fin clipping | PGP_1 | X | 5 | AGGCTGA | PGP-C57G8ANXX_5_fastq.gz |
| 68 | FAR-F_07 | 22/10 2013 | Perca_fluviatilis | Fårup Sø | FAR-F | 67 | P45627 | 41858 | Adult | Fresh | Denmark | Fin clipping | PGP_1 | X | 5 | GCTGAGA | PGP-C57G8ANXX_5_fastq.gz |
| 69 | FAR-F_08 | 22/10 2013 | Perca_fluviatilis | Fårup Sø | FAR-F | 68 | P45628 | 41858 | Adult | Fresh | Denmark | Fin clipping | PGP_1 | X | 5 | GGCTATA | PGP-C57G8ANXX_5_fastq.gz |
| 70 | FAR-F_09 | 22/10 2013 | Perca_fluviatilis | Fårup Sø | FAR-F | 69 | P45629 | 41858 | Adult | Fresh | Denmark | Fin clipping | PGP_1 | X | 5 | CCTAATCG | PGP-C57G8ANXX_5_fastq.gz |
| 71 | FAR-F_10 | 22/10 2013 | Perca_fluviatilis | Fårup Sø | FAR-F | 70 | P45630 | 41858 | Adult | Fresh | Denmark | Fin clipping | PGP_1 | X | 5 | CTCAAGCG | PGP-C57G8ANXX_5_fastq.gz |
| 72 | KET-B_01 | 06/11 2013 | Perca_fluviatilis | Ketting Nor | KET-B | 71 | P45681 | 41858 | Juvenile | Brackish | Denmark | Fin clipping | PGP_1 | X | 5 | CGCCATCG | PGP-C57G8ANXX_5_fastq.gz |
| 73 | KET-B_02 | 06/11 2013 | Perca_fluviatilis | Ketting Nor | KET-B | 72 | P45682 | 41858 | Juvenile | Brackish | Denmark | Fin clipping | PGP_1 | X | 5 | ACCTAG | PGP-C57G8ANXX_5_fastq.gz |
| 74 | KET-B_03 | 06/11 2013 | Perca_fluviatilis | Ketting Nor | KET-B | 73 | P45683 | 41858 | Juvenile | Brackish | Denmark | Fin clipping | PGP_1 | X | 5 | GGACTAA | PGP-C57G8ANXX_5_fastq.gz |
| 75 | KET-B_04 | 06/11 2013 | Perca_fluviatilis | Ketting Nor | KET-B | 74 | P45684 | 41858 | Juvenile | Brackish | Denmark | Fin clipping | PGP_1 | X | 5 | GCAATAG | PGP-C57G8ANXX_5_fastq.gz |
| 76 | KET-B_05 | 06/11 2013 | Perca_fluviatilis | Ketting Nor | KET-B | 75 | P45685 | 41858 | Juvenile | Brackish | Denmark | Fin clipping | PGP_1 | X | 5 | AACGTAG | PGP-C57G8ANXX_5_fastq.gz |
| 77 | KET-B_06 | 06/11 2013 | Perca_fluviatilis | Ketting Nor | KET-B | 76 | P45686 | 41858 | Juvenile | Brackish | Denmark | Fin clipping | PGP_1 | X | 5 | CAGGTCG | PGP-C57G8ANXX_5_fastq.gz |
| 78 | KET-B_07 | 06/11 2013 | Perca_fluviatilis | Ketting Nor | KET-B | 77 | P45687 | 41858 | Juvenile | Brackish | Denmark | Fin clipping | PGP_1 | X | 5 | CGCTTCCA | PGP-C57G8ANXX_5_fastq.gz |
| 79 | KET-B_08 | 06/11 2013 | Perca_fluviatilis | Ketting Nor | KET-B | 78 | P45688 | 41858 | Juvenile | Brackish | Denmark | Fin clipping | PGP_1 | X | 5 | TCTTGCAA | PGP-C57G8ANXX_5_fastq.gz |
| 80 | KET-B_09 | 06/11 2013 | Perca_fluviatilis | Ketting Nor | KET-B | 79 | P45689 | 41858 | Juvenile | Brackish | Denmark | Fin clipping | PGP_1 | X | 5 | TCTACGAA | PGP-C57G8ANXX_5_fastq.gz |
| 81 | KET-B_10 | 06/11 2013 | Perca_fluviatilis | Ketting Nor | KET-B | 80 | P45690 | 41858 | Juvenile | Brackish | Denmark | Fin clipping | PGP_1 | X | 5 | CGTCGA | PGP-C57G8ANXX_5_fastq.gz |
| 82 | KET-B_11 | 06/11 2013 | Perca_fluviatilis | Ketting Nor | KET-B | 81 | P45691 | 41858 | Juvenile | Brackish | Denmark | Fin clipping | PGP_1 | X | 5 | AAGTACG | PGP-C57G8ANXX_5_fastq.gz |
| 83 | KET-B_12 | 06/11 2013 | Perca_fluviatilis | Ketting Nor | KET-B | 82 | P45692 | 41858 | Juvenile | Brackish | Denmark | Fin clipping | PGP_1 | X | 5 | CATCAGA | PGP-C57G8ANXX_5_fastq.gz |
| 84 | KET-B_13 | 06/11 2013 | Perca_fluviatilis | Ketting Nor | KET-B | 83 | P45693 | 41858 | Adult | Brackish | Denmark | Fin clipping | PGP_1 | X | 5 | CGATCAA | PGP-C57G8ANXX_5_fastq.gz |
| 85 | SJA-F_01 | 16/12 2013 | Perca_fluviatilis | Sjælsø | SJA-F | 86 | P45636 | 41858 | Adult | Fresh | Denmark | Fin clipping | PGP_1 | X | 5 | CGCACTCA | PGP-C57G8ANXX_5_fastq.gz |
| 86 | SJA-F_02 | 16/12 2013 | Perca_fluviatilis | Sjælsø | SJA-F | 87 | P45637 | 13/02 2014 | Adult | Fresh | Denmark | Fin clipping | PGP_1 | X | 5 | ATGGCGCG | PGP-C57G8ANXX_5_fastq.gz |
| 87 | SJA-F_03 | 16/12 2013 | Perca_fluviatilis | Sjælsø | SJA-F | 88 | P45638 | 41858 | Adult | Fresh | Denmark | Fin clipping | PGP_1 | X | 5 | GAGGCTCG | PGP-C57G8ANXX_5_fastq.gz |
| 88 | SJA-F_04 | 16/12 2013 | Perca_fluviatilis | Sjælsø | SJA-F | 89 | P45639 | 41858 | Juvenile | Fresh | Denmark | Fin clipping | PGP_1 | X | 5 | ATGTGACG | PGP-C57G8ANXX_5_fastq.gz |
| 89 | SJA-F_05 | 16/12 2013 | Perca_fluviatilis | Sjælsø | SJA-F | 90 | P45640 | 41858 | Adult | Fresh | Denmark | Fin clipping | PGP_1 | X | 5 | ATGACG | PGP-C57G8ANXX_5_fastq.gz |
| 90 | SJA-F_06 | 16/12 2013 | Perca_fluviatilis | Sjælsø | SJA-F | 91 | P45653 | 41858 | Adult | Fresh | Denmark | Fin clipping | PGP_1 | X | 5 | TCCGCAG | PGP-C57G8ANXX_5_fastq.gz |
| 91 | SJA-F_07 | 16/12 2013 | Perca_fluviatilis | Sjælsø | SJA-F | 92 | P45654 | 41858 | Adult | Fresh | Denmark | Fin clipping | PGP_1 | X | 5 | ATCGGCG | PGP-C57G8ANXX_5_fastq.gz |
| 92 | SJA-F_08 | 16/12 2013 | Perca_fluviatilis | Sjælsø | SJA-F | 93 | P45655 | 41858 | Adult | Fresh | Denmark | Fin clipping | PGP_1 | X | 5 | ACTCTCG | PGP-C57G8ANXX_5_fastq.gz |
| 93 | SJA-F_09 | 16/12 2013 | Perca_fluviatilis | Sjælsø | SJA-F | 94 | P45656 | 41858 | Juvenile | Fresh | Denmark | Fin clipping | PGP_1 | X | 5 | GACATCCA | PGP-C57G8ANXX_5_fastq.gz |
| 94 | SJA-F_10 | 16/12 2013 | Perca_fluviatilis | Sjælsø | SJA-F | 95 | P45657 | 41858 | Juvenile | Fresh | Denmark | Fin clipping | PGP_1 | X | 5 | TAAGCTCA | PGP-C57G8ANXX_5_fastq.gz |
| 95 | SJA-F_11 | 16/12 2013 | Perca_fluviatilis | Sjælsø | SJA-F | 96 | P45658 | 41858 | Juvenile | Fresh | Denmark | Fin clipping | PGP_1 | X | 5 | AGATCGAA | PGP-C57G8ANXX_5_fastq.gz |
| 96 | SJA-F_12 | 16/12 2013 | Perca_fluviatilis | Sjælsø | SJA-F | 97 | P45659 | 41858 | Juvenile | Fresh | Denmark | Fin clipping | PGP_1 | X | 5 | GAAGTACA | PGP-C57G8ANXX_5_fastq.gz |
| 97 | TYB-F_01 | 20/11 2013 | Perca_fluviatilis | Bavelse sø | TYB-F | 84 | P45634 | 2/13/2014 | Adult | Fresh | Denmark | Fin clipping | PGP_2 | X | 6 | TTGA | PGP-C57G8ANXX_6_fastq.gz |
| 98 | TYB-F_02 | 20/11 2014 | Perca_fluviatilis | Bavelse sø | TYB-F | 85 | P45635 | 2/13/2014 | Adult | Fresh | Denmark | Fin clipping | PGP_2 | X | 6 | CTAGAA | PGP-C57G8ANXX_6_fastq.gz |
| 99 | ISH-B_01 | 08/02 2014 | Perca_fluviatilis | Ishøj Havn | ISH-B | 99 | p45702 | | Adult | Brackish | Denmark | Fin clipping | PGP_2 | X | 6 | CGTAGGA | PGP-C57G8ANXX_6_fastq.gz |
| 100 | ISH-B_02 | 08/02 2014 | Perca_fluviatilis | Ishøj Havn | ISH-B | 101 | p45704 | | Adult | Brackish | Denmark | Fin clipping | PGP_2 | X | 6 | TGGTCAG | PGP-C57G8ANXX_6_fastq.gz |
| 101 | ISH-B_03 | 08/02 2014 | Perca_fluviatilis | Ishøj Havn | ISH-B | 102 | p45705 | | Adult | Brackish | Denmark | Fin clipping | PGP_2 | X | 6 | TTAACGA | PGP-C57G8ANXX_6_fastq.gz |
| 102 | ISH-B_04 | 08/02 2014 | Perca_fluviatilis | Ishøj Havn | ISH-B | 103 | p45706 | | Adult | Brackish | Denmark | Fin clipping | PGP_2 | X | 6 | ACTCGACA | PGP-C57G8ANXX_6_fastq.gz |
| 103 | ISH-B_05 | 08/02 2014 | Perca_fluviatilis | Ishøj Havn | ISH-B | 104 | p45707 | | Adult | Brackish | Denmark | Fin clipping | PGP_2 | X | 6 | GCCATACG | PGP-C57G8ANXX_6_fastq.gz |
| 104 | ISH-B_06 | 08/02 2014 | Perca_fluviatilis | Ishøj Havn | ISH-B | 105 | p45708 | | Adult | Brackish | Denmark | Fin clipping | PGP_2 | X | 6 | CTCCGACG | PGP-C57G8ANXX_6_fastq.gz |
| 105 | ISH-B_07 | 08/02 2014 | Perca_fluviatilis | Ishøj Havn | ISH-B | 106 | p45709 | | Adult | Brackish | Denmark | Fin clipping | PGP_2 | X | 6 | GGTA | PGP-C57G8ANXX_6_fastq.gz |
| 106 | ISH-B_08 | 08/02 2014 | Perca_fluviatilis | Ishøj Havn | ISH-B | 107 | p45710 | | Adult | Brackish | Denmark | Fin clipping | PGP_2 | X | 6 | GATACG | PGP-C57G8ANXX_6_fastq.gz |
| 107 | ISH-B_09 | 08/02 2014 | Perca_fluviatilis | Ishøj Havn | ISH-B | 108 | p45711 | | Adult | Brackish | Denmark | Fin clipping | PGP_2 | X | 6 | GATAGCG | PGP-C57G8ANXX_6_fastq.gz |
| 108 | ISH-B_10 | 08/02 2014 | Perca_fluviatilis | Ishøj Havn | ISH-B | 109 | p45712 | | Adult | Brackish | Denmark | Fin clipping | PGP_2 | X | 6 | GACGTGA | PGP-C57G8ANXX_6_fastq.gz |
| 109 | ISH-B_11 | 08/02 2014 | Perca_fluviatilis | Ishøj Havn | ISH-B | 110 | p45713 | | Adult | Brackish | Denmark | Fin clipping | PGP_2 | X | 6 | TAGCGCG | PGP-C57G8ANXX_6_fastq.gz |
| 110 | ISH-B_12 | 08/02 2014 | Perca_fluviatilis | Ishøj Havn | ISH-B | 111 | p45714 | | Adult | Brackish | Denmark | Fin clipping | PGP_2 | X | 6 | TGAGAGCA | PGP-C57G8ANXX_6_fastq.gz |
| 111 | ISH-B_13 | 08/02 2014 | Perca_fluviatilis | Ishøj Havn | ISH-B | 112 | p45715 | | Adult | Brackish | Denmark | Fin clipping | PGP_2 | X | 6 | ATTCAGCA | PGP-C57G8ANXX_6_fastq.gz |
| 112 | ISH-B_14 | 08/02 2014 | Perca_fluviatilis | Ishøj Havn | ISH-B | 113 | p45716 | | Adult | Brackish | Denmark | Fin clipping | PGP_2 | X | 6 | TCTGATAA | PGP-C57G8ANXX_6_fastq.gz |
| 113 | ISH-B_15 | 08/02 2014 | Perca_fluviatilis | Ishøj Havn | ISH-B | 114 | p45717 | | Adult | Brackish | Denmark | Fin clipping | PGP_2 | X | 6 | TCTGG | PGP-C57G8ANXX_6_fastq.gz |
| 114 | ISH-B_16 | 08/02 2014 | Perca_fluviatilis | Ishøj Havn | ISH-B | 115 | p45718 | | Adult | Brackish | Denmark | Fin clipping | PGP_2 | X | 6 | TCCGAA | PGP-C57G8ANXX_6_fastq.gz |
| 115 | ISH-B_17 | 08/02 2014 | Perca_fluviatilis | Ishøj Havn | ISH-B | 116 | Blank | | Adult | Brackish | Denmark | Fin clipping | PGP_2 | X | 6 | ACCTCGA | PGP-C57G8ANXX_6_fastq.gz |
| 116 | ISH-B_18 | 08/02 2014 | Perca_fluviatilis | Ishøj Havn | ISH-B | 117 | Blank | | Adult | Brackish | Denmark | Fin clipping | PGP_2 | X | 6 | CTAAGAA | PGP-C57G8ANXX_6_fastq.gz |

| | | | | | | | | | | | | | | | | | |
|---|---|---|---|---|---|---|---|---|---|---|---|---|---|---|---|---|---|
| 117 | RAN-B_13 | 04/03 2014 | Perca_fluviatilis | Randers Fjord | RAN-B | 118 | p45557 | | Juvenile | Brackish | Denmark | Fin clipping | PGP_2 | X | 6 | GCGTTAA | PGP-C57G8ANXX_6_fastq.gz |
| 118 | RAN-B_14 | 04/03 2014 | Perca_fluviatilis | Randers Fjord | RAN-B | 119 | p45558 | | Juvenile | Brackish | Denmark | Fin clipping | PGP_2 | X | 6 | GTGTGCCA | PGP-C57G8ANXX_6_fastq.gz |
| 119 | TYB-F_03 | 09/03 2014 | Perca_fluviatilis | Tystrup-Bavelse Sø | TYB-F | 120 | P45720 | | Adult | Fresh | Denmark | Fin clipping | PGP_2 | X | 6 | TAGTGCCG | PGP-C57G8ANXX_6_fastq.gz |
| 120 | TYB-F_04 | 09/03 2014 | Perca_fluviatilis | Tystrup-Bavelse Sø | TYB-F | 121 | P45721 | | Adult | Fresh | Denmark | Fin clipping | PGP_2 | X | 6 | AGGATCCG | PGP-C57G8ANXX_6_fastq.gz |
| 121 | TYB-F_05 | 09/03 2014 | Perca_fluviatilis | Tystrup-Bavelse Sø | TYB-F | 122 | P45722 | | Adult | Fresh | Denmark | Fin clipping | PGP_2 | X | 6 | CGGCA | PGP-C57G8ANXX_6_fastq.gz |
| 122 | TYB-F_06 | 09/03 2014 | Perca_fluviatilis | Tystrup-Bavelse Sø | TYB-F | 123 | P45723 | | Adult | Fresh | Denmark | Fin clipping | PGP_2 | X | 6 | CAGTGA | PGP-C57G8ANXX_6_fastq.gz |
| 123 | TYB-F_07 | 09/03 2014 | Perca_fluviatilis | Tystrup-Bavelse Sø | TYB-F | 124 | P45724 | | Adult | Fresh | Denmark | Fin clipping | PGP_2 | X | 6 | TTGCTAA | PGP-C57G8ANXX_6_fastq.gz |
| 124 | TYB-F_08 | 09/03 2014 | Perca_fluviatilis | Tystrup-Bavelse Sø | TYB-F | 125 | P45725 | | Adult | Fresh | Denmark | Fin clipping | PGP_2 | X | 6 | GGTCACG | PGP-C57G8ANXX_6_fastq.gz |
| 125 | TYB-F_09 | 09/03 2014 | Perca_fluviatilis | Tystrup-Bavelse Sø | TYB-F | 126 | P45726 | | Adult | Fresh | Denmark | Fin clipping | PGP_2 | X | 6 | CTCAAGA | PGP-C57G8ANXX_6_fastq.gz |
| 126 | TYB-F_10 | 09/03 2014 | Perca_fluviatilis | Tystrup-Bavelse Sø | TYB-F | 127 | P45727 | | Adult | Fresh | Denmark | Fin clipping | PGP_2 | X | 6 | AAGACTCA | PGP-C57G8ANXX_6_fastq.gz |
| 127 | PGP_2-Blank | PGP_2-Blank | Na | Na | Na | Na | Na | Na | Na | Na | Na | Na | PGP_2 | X | 6 | GGCTATCA | PGP-C57G8ANXX_6_fastq.gz |
| 128 | POL-BF_01 | | Perca_fluviatilis | Pøle Å | POL-BF | 128 | p45739 | | Juvenile | Fresh | Denmark | Fin clipping | PGP_2 | X | 6 | GAATACAA | PGP-C57G8ANXX_6_fastq.gz |
| 129 | POL-BF_02 | | Perca_fluviatilis | Pøle Å | POL-BF | 129 | p45740 | | Juvenile | Fresh | | | PGP_2 | X | 6 | GAAGA | PGP-C57G8ANXX_6_fastq.gz |
| 130 | POL-BF_03 | | Perca_fluviatilis | Pøle Å | POL-BF | 130 | p45741 | | Juvenile | Fresh | Denmark | Fin clipping | PGP_2 | X | 6 | AGTCCG | PGP-C57G8ANXX_6_fastq.gz |
| 131 | POL-BF_04 | | Perca_fluviatilis | Pøle Å | POL-BF | 131 | p45742 | | Juvenile | Fresh | Denmark | Fin clipping | PGP_2 | X | 6 | TGAGGCG | PGP-C57G8ANXX_6_fastq.gz |
| 132 | POL-BF_05 | | Perca_fluviatilis | Pøle Å | POL-BF | 132 | p45743 | | Juvenile | Fresh | Denmark | Fin clipping | PGP_2 | X | 6 | TCGTAGA | PGP-C57G8ANXX_6_fastq.gz |
| 133 | POL-BF_06 | | Perca_fluviatilis | Pøle Å | POL-BF | 133 | p45744 | | Juvenile | Fresh | Denmark | Fin clipping | PGP_2 | X | 6 | CGTGGAG | PGP-C57G8ANXX_6_fastq.gz |
| 134 | POL-BF_07 | | Perca_fluviatilis | Pøle Å | POL-BF | 134 | p45745 | | Juvenile | Fresh | Denmark | Fin clipping | PGP_2 | X | 6 | CCACTACG | PGP-C57G8ANXX_6_fastq.gz |
| 135 | POL-BF_08 | | Perca_fluviatilis | Pøle Å | POL-BF | 135 | p45746 | | Juvenile | Fresh | Denmark | Fin clipping | PGP_2 | X | 6 | CCTGCACG | PGP-C57G8ANXX_6_fastq.gz |
| 136 | POL-BF_09 | | Perca_fluviatilis | Pøle Å | POL-BF | 136 | p45747 | | Juvenile | Fresh | Denmark | Fin clipping | PGP_2 | X | 6 | CTGACACG | PGP-C57G8ANXX_6_fastq.gz |
| 137 | POL-BF_10 | | Perca_fluviatilis | Pøle Å | POL-BF | 137 | p45748 | | Juvenile | Fresh | Denmark | Fin clipping | PGP_2 | X | 6 | ACGAG | PGP-C57G8ANXX_6_fastq.gz |
| 138 | POL-BF_11 | | Perca_fluviatilis | Pøle Å | POL-BF | 138 | p45749 | | Juvenile | Fresh | Denmark | Fin clipping | PGP_2 | X | 6 | GCATGG | PGP-C57G8ANXX_6_fastq.gz |
| 139 | POL-BF_12 | | Perca_fluviatilis | Pøle Å | POL-BF | 139 | p45750 | | Juvenile | Fresh | Denmark | Fin clipping | PGP_2 | X | 6 | GACTAGG | PGP-C57G8ANXX_6_fastq.gz |
| 140 | POL-BF_13 | | Perca_fluviatilis | Pøle Å | POL-BF | 140 | p45751 | | Juvenile | Fresh | Denmark | Fin clipping | PGP_2 | X | 6 | AGCTCCG | PGP-C57G8ANXX_6_fastq.gz |
| 141 | POL-BF_14 | | Perca_fluviatilis | Pøle Å | POL-BF | 141 | p45752 | | Juvenile | Fresh | Denmark | Fin clipping | PGP_2 | X | 6 | GTACGCG | PGP-C57G8ANXX_6_fastq.gz |
| 142 | POL-BF_15 | | Perca_fluviatilis | Pøle Å | POL-BF | 142 | p45753 | | Juvenile | Fresh | Denmark | Fin clipping | PGP_2 | X | 6 | GGTTCGCA | PGP-C57G8ANXX_6_fastq.gz |
| 143 | POL-BF_16 | | Perca_fluviatilis | Pøle Å | POL-BF | 143 | p45754 | | Juvenile | Fresh | Denmark | Fin clipping | PGP_2 | X | 6 | TTAAGCAA | PGP-C57G8ANXX_6_fastq.gz |
| 144 | POL-BF_17 | | Perca_fluviatilis | Pøle Å | POL-BF | 144 | p45755 | | Juvenile | Fresh | Denmark | Fin clipping | PGP_2 | X | 6 | TCCGTGAA | PGP-C57G8ANXX_6_fastq.gz |
| 145 | POL-BF_18 | | Perca_fluviatilis | Pøle Å | POL-BF | 145 | p45756 | | Juvenile | Fresh | Denmark | Fin clipping | PGP_2 | X | 6 | TTCTA | PGP-C57G8ANXX_6_fastq.gz |
| 146 | RAN-B_15 | 41796 | Perca_fluviatilis | Randers Fjord | RAN-B | 146 | p45593 | | Adult | Brackish | Denmark | Fin clipping | PGP_2 | X | 6 | GTCCAA | PGP-C57G8ANXX_6_fastq.gz |
| 147 | RAN-B_16 | 41796 | Perca_fluviatilis | Randers Fjord | RAN-B | 147 | p45594 | | Adult | Brackish | Denmark | Fin clipping | PGP_2 | X | 6 | ATAGCAA | PGP-C57G8ANXX_6_fastq.gz |
| 148 | RAN-B_17 | 41796 | Perca_fluviatilis | Randers Fjord | RAN-B | 148 | p45595 | | Adult | Brackish | Denmark | Fin clipping | PGP_2 | X | 6 | TAACGAA | PGP-C57G8ANXX_6_fastq.gz |
| 149 | RAN-B_18 | 41796 | Perca_fluviatilis | Randers Fjord | RAN-B | 149 | p45596 | | Adult | Brackish | Denmark | Fin clipping | PGP_2 | X | 6 | AACGCTA | PGP-C57G8ANXX_6_fastq.gz |
| 150 | RAN-B_19 | 41796 | Perca_fluviatilis | Randers Fjord | RAN-B | 150 | p45597 | | Adult | Brackish | Denmark | Fin clipping | PGP_2 | X | 6 | ATCGACCG | PGP-C57G8ANXX_6_fastq.gz |
| 151 | RAN-B_20 | 41796 | Perca_fluviatilis | Randers Fjord | RAN-B | 151 | p45598 | | Adult | Brackish | Denmark | Fin clipping | PGP_2 | X | 6 | GAGCTGCA | PGP-C57G8ANXX_6_fastq.gz |
| 152 | RAN-B_21 | 41796 | Perca_fluviatilis | Randers Fjord | RAN-B | 152 | p45599 | | Adult | Brackish | Denmark | Fin clipping | PGP_2 | X | 6 | GATCGTCG | PGP-C57G8ANXX_6_fastq.gz |
| 153 | RAN-B_22 | 41796 | Perca_fluviatilis | Randers Fjord | RAN-B | 153 | p45600 | | Adult | Brackish | Denmark | Fin clipping | PGP_2 | X | 6 | AGTCG | PGP-C57G8ANXX_6_fastq.gz |
| 154 | RAN-B_23 | 41796 | Perca_fluviatilis | Randers Fjord | RAN-B | 154 | p45758 | | Adult | Brackish | Denmark | Fin clipping | PGP_2 | X | 6 | CAGGTG | PGP-C57G8ANXX_6_fastq.gz |
| 155 | RAN-B_24 | 41796 | Perca_fluviatilis | Randers Fjord | RAN-B | 155 | p45759 | | Adult | Brackish | Denmark | Fin clipping | PGP_2 | X | 6 | CCTAACG | PGP-C57G8ANXX_6_fastq.gz |
| 156 | RAN-B_25 | 41796 | Perca_fluviatilis | Randers Fjord | RAN-B | 156 | p45760 | | Adult | Brackish | Denmark | Fin clipping | PGP_2 | X | 6 | CTGACCG | PGP-C57G8ANXX_6_fastq.gz |
| 157 | RAN-B_26 | 41796 | Perca_fluviatilis | Randers Fjord | RAN-B | 157 | p45761 | | Adult | Brackish | Denmark | Fin clipping | PGP_2 | X | 6 | TCTATGG | PGP-C57G8ANXX_6_fastq.gz |
| 158 | RAN-B_27 | 41796 | Perca_fluviatilis | Randers Fjord | RAN-B | 158 | p45762 | | Adult | Brackish | Denmark | Fin clipping | PGP_2 | X | 6 | TAACGACA | PGP-C57G8ANXX_6_fastq.gz |
| 159 | RAN-B_28 | 41796 | Perca_fluviatilis | Randers Fjord | RAN-B | 159 | p45763 | | Adult | Brackish | Denmark | Fin clipping | PGP_2 | X | 6 | AGACTAAG | PGP-C57G8ANXX_6_fastq.gz |
| 160 | RAN-B_29 | 41796 | Perca_fluviatilis | Randers Fjord | RAN-B | 160 | p45764 | | Adult | Brackish | Denmark | Fin clipping | PGP_2 | X | 6 | AGAGTCAA | PGP-C57G8ANXX_6_fastq.gz |
| 161 | RAN-B_30 | 41796 | Perca_fluviatilis | Randers Fjord | RAN-B | 161 | p45765 | | Adult | Brackish | Denmark | Fin clipping | PGP_2 | X | 6 | TACAGA | PGP-C57G8ANXX_6_fastq.gz |
| 162 | TAN-F_13 | 41796 | Perca_fluviatilis | Tange Sø | TAN-F | 162 | p45573 | | Juvenile | Fresh | Denmark | Fin clipping | PGP_2 | X | 6 | TCACGA | PGP-C57G8ANXX_6_fastq.gz |
| 163 | TAN-F_14 | 41796 | Perca_fluviatilis | Tange Sø | TAN-F | 163 | p45574 | | Juvenile | Fresh | Denmark | Fin clipping | PGP_2 | X | 6 | AGGCTGA | PGP-C57G8ANXX_6_fastq.gz |
| 164 | TAN-F_15 | 41796 | Perca_fluviatilis | Tange Sø | TAN-F | 164 | p45575 | | Juvenile | Fresh | Denmark | Fin clipping | PGP_2 | X | 6 | GCTGAGA | PGP-C57G8ANXX_6_fastq.gz |
| 165 | TAN-F_16 | 41796 | Perca_fluviatilis | Tange Sø | TAN-F | 165 | p45576 | | Juvenile | Fresh | Denmark | Fin clipping | PGP_2 | X | 6 | GGCTATA | PGP-C57G8ANXX_6_fastq.gz |
| 166 | TAN-F_17 | 41796 | Perca_fluviatilis | Tange Sø | TAN-F | 166 | p45577 | | Juvenile | Fresh | Denmark | Fin clipping | PGP_2 | X | 6 | CCTAATCG | PGP-C57G8ANXX_6_fastq.gz |
| 167 | TAN-F_18 | 41796 | Perca_fluviatilis | Tange Sø | TAN-F | 167 | p45578 | | Juvenile | Fresh | Denmark | Fin clipping | PGP_2 | X | 6 | CTCAAGCG | PGP-C57G8ANXX_6_fastq.gz |
| 168 | TAN-F_19 | 41796 | Perca_fluviatilis | Tange Sø | TAN-F | 168 | p45579 | | Juvenile | Fresh | Denmark | Fin clipping | PGP_2 | X | 6 | CGCCATCG | PGP-C57G8ANXX_6_fastq.gz |
| 169 | TYB-F_11 | 41888 | Perca_fluviatilis | Tystrup-Bavelse Sø | TYB-F | 169 | p45766 | | Adult | Fresh | Denmark | Fin clipping | PGP_2 | X | 6 | ACCTAG | PGP-C57G8ANXX_6_fastq.gz |
| 170 | TYB-F_12 | 41888 | Perca_fluviatilis | Tystrup-Bavelse Sø | TYB-F | 170 | p45767 | | Adult | Fresh | Denmark | Fin clipping | PGP_2 | X | 6 | GGACTAA | PGP-C57G8ANXX_6_fastq.gz |
| 171 | TYB-F_13 | 41888 | Perca_fluviatilis | Tystrup-Bavelse Sø | TYB-F | 171 | p45768 | | Adult | Fresh | Denmark | Fin clipping | PGP_2 | X | 6 | GCAATAG | PGP-C57G8ANXX_6_fastq.gz |
| 172 | KAR-B_13 | 43204 | Perca_fluviatilis | Karrebæk Fjord | KAR-B | 172 | p45728 | | Adult | Brackish | Denmark | Fin clipping | PGP_2 | X | 6 | AACGTAG | PGP-C57G8ANXX_6_fastq.gz |
| 173 | KAR-B_14 | 43204 | Perca_fluviatilis | Karrebæk Fjord | KAR-B | 173 | p45729 | | Adult | Brackish | Denmark | Fin clipping | PGP_2 | X | 6 | CAGGTCG | PGP-C57G8ANXX_6_fastq.gz |
| 174 | KAR-B_15 | 43204 | Perca_fluviatilis | Karrebæk Fjord | KAR-B | 174 | p45730 | | Adult | Brackish | Denmark | Fin clipping | PGP_2 | X | 6 | CGCTTCCA | PGP-C57G8ANXX_6_fastq.gz |
| 175 | KAR-B_16 | 43204 | Perca_fluviatilis | Karrebæk Fjord | KAR-B | 175 | p45731 | | Adult | Brackish | Denmark | Fin clipping | PGP_2 | X | 6 | TCTTGCAA | PGP-C57G8ANXX_6_fastq.gz |

| | | | | | | | | | | | | | | | | |
|---|---|---|---|---|---|---|---|---|---|---|---|---|---|---|---|---|
| 176 | KAR-B_17 | 43204 | Perca_fluviatilis | Karrebæk Fjord | KAR-B | 176 | p45732 | Adult | Brackish | Denmark | Fin clipping | PGP_2 | X | 6 | TCTACGAA | PGP-C57G8ANXX_6_fastq.gz |
| 177 | KAR-B_18 | 43204 | Perca_fluviatilis | Karrebæk Fjord | KAR-B | 177 | p45733 | Adult | Brackish | Denmark | Fin clipping | PGP_2 | X | 6 | CGTCGA | PGP-C57G8ANXX_6_fastq.gz |
| 178 | KAR-B_19 | 43204 | Perca_fluviatilis | Karrebæk Fjord | KAR-B | 178 | p45734 | Adult | Brackish | Denmark | Fin clipping | PGP_2 | X | 6 | AAGTACG | PGP-C57G8ANXX_6_fastq.gz |
| 179 | KAR-B_20 | 43204 | Perca_fluviatilis | Karrebæk Fjord | KAR-B | 179 | p45735 | Adult | Brackish | Denmark | Fin clipping | PGP_2 | X | 6 | CATCAGA | PGP-C57G8ANXX_6_fastq.gz |
| 180 | SON-F_01 | 43326 | Perca_fluviatilis | Sønder Sø | SON-F | 180 | Na | Adult | Fresh | Denmark | Fin clipping | PGP_2 | X | 6 | CGATCAA | PGP-C57G8ANXX_6_fastq.gz |
| 181 | SON-F_02 | 43326 | Perca_fluviatilis | Sønder Sø | SON-F | 181 | Na | Adult | Fresh | Denmark | Fin clipping | PGP_2 | X | 6 | CGCACTCA | PGP-C57G8ANXX_6_fastq.gz |
| 182 | SON-F_03 | 43326 | Perca_fluviatilis | Sønder Sø | SON-F | 182 | Na | Adult | Fresh | Denmark | Fin clipping | PGP_2 | X | 6 | ATGGCGCG | PGP-C57G8ANXX_6_fastq.gz |
| 183 | SON-F_04 | 43326 | Perca_fluviatilis | Sønder Sø | SON-F | 183 | Na | Adult | Fresh | Denmark | Fin clipping | PGP_2 | X | 6 | GAGGCTCG | PGP-C57G8ANXX_6_fastq.gz |
| 184 | SON-F_05 | 43326 | Perca_fluviatilis | Sønder Sø | SON-F | 184 | Na | Adult | Fresh | Denmark | Fin clipping | PGP_2 | X | 6 | ATGTGACG | PGP-C57G8ANXX_6_fastq.gz |
| 185 | SON-F_06 | 43326 | Perca_fluviatilis | Sønder Sø | SON-F | 185 | Na | Adult | Fresh | Denmark | Fin clipping | PGP_2 | X | 6 | ATGACG | PGP-C57G8ANXX_6_fastq.gz |
| 186 | SON-F_07 | 43326 | Perca_fluviatilis | Sønder Sø | SON-F | 186 | Na | Adult | Fresh | Denmark | Fin clipping | PGP_2 | X | 6 | TCCGCAG | PGP-C57G8ANXX_6_fastq.gz |
| 187 | SON-F_08 | 43326 | Perca_fluviatilis | Sønder Sø | SON-F | 187 | Na | Adult | Fresh | Denmark | Fin clipping | PGP_2 | X | 6 | ATCGGCG | PGP-C57G8ANXX_6_fastq.gz |
| 188 | SON-F_09 | 43326 | Perca_fluviatilis | Sønder Sø | SON-F | 188 | Na | Adult | Fresh | Denmark | Fin clipping | PGP_2 | X | 6 | ACTCTCG | PGP-C57G8ANXX_6_fastq.gz |
| 189 | SON-F_10 | 43326 | Perca_fluviatilis | Sønder Sø | SON-F | 189 | Na | Adult | Fresh | Denmark | Fin clipping | PGP_2 | X | 6 | GACATCCA | PGP-C57G8ANXX_6_fastq.gz |
| 190 | SON-F_11 | 43326 | Perca_fluviatilis | Sønder Sø | SON-F | 190 | Na | Adult | Fresh | Denmark | Fin clipping | PGP_2 | X | 6 | TAAGCTCA | PGP-C57G8ANXX_6_fastq.gz |
| 191 | SON-F_12 | 43326 | Perca_fluviatilis | Sønder Sø | SON-F | 191 | Na | Adult | Fresh | Denmark | Fin clipping | PGP_2 | X | 6 | AGATCGAA | PGP-C57G8ANXX_6_fastq.gz |
| 192 | SON-F_13 | 43326 | Perca_fluviatilis | Sønder Sø | SON-F | 192 | Na | Adult | Fresh | Denmark | Fin clipping | PGP_2 | X | 6 | GAAGTACA | PGP-C57G8ANXX_6_fastq.gz |
