## Supplementary material for "Genomic differentiation among European perch in the western Baltic Sea reflects colonisation history and local adaptation": Table S2

Table S2: mean generation time of eight populations of European perch, originating from brackish water (BW) and fresh water (FW). The mean asymptotic length (L∞) and growth factor (K) from von Bertalanfy growht curve fits in Christensen et al (2021) was used to calculate mean length of maximum yield (Lopt) (Froese and Bihnolan 2000), assuming that the corresponding age is equivalent to generation time, as suggested by Beverton (1992). Calculations we performed according to: TL = TL∞ * (1-exp(-k*t)) + 2

| **Country** | **Locality** | **Habitat** | **L∞** | **K** | **Lopt** | **Gen time** |
|---|---|---|---|---|---|---|
| Denmark | Flintinge Å | BW | 33,3 | 0,4 | 20,5 | 1,8 |
| Ireland | Lough Rea | FW | 42,9 | 0,2 | 26,7 | 4,8 |
| The Netherlands | Lerwersmeer | FW | 45,7 | 0,3 | 28,5 | 3,3 |
| Ireland | Loguh Glore | FW | 13,7 | 0,3 | 8,1 | 2,3 |
| The Netherlands | Lake Ijssel | FW | 44,1 | 0,2 | 27,5 | 4,3 |
| Germany | Bodensee | FW | 61,3 | 0,2 | 38,8 | 5,8 |
| Scotland | Dubh Lochan | FW | 39,6 | 0,2 | 24,6 | 3,4 |
| Demark | Tryggevælde Å | BW | 44,7 | 0,3 | 27,9 | 3,4 |
| Denmark | St vejle Å | BW | 41,3 | 0,2 | 25,7 | 4,2 |
| | | Median | 42,9 | 0,2 | 26,7 | 3,4 |
