## Supplementary figures and images for "Genomic differentiation among European perch in the western Baltic Sea reflects colonisation history and local adaptation"

### Figure S1

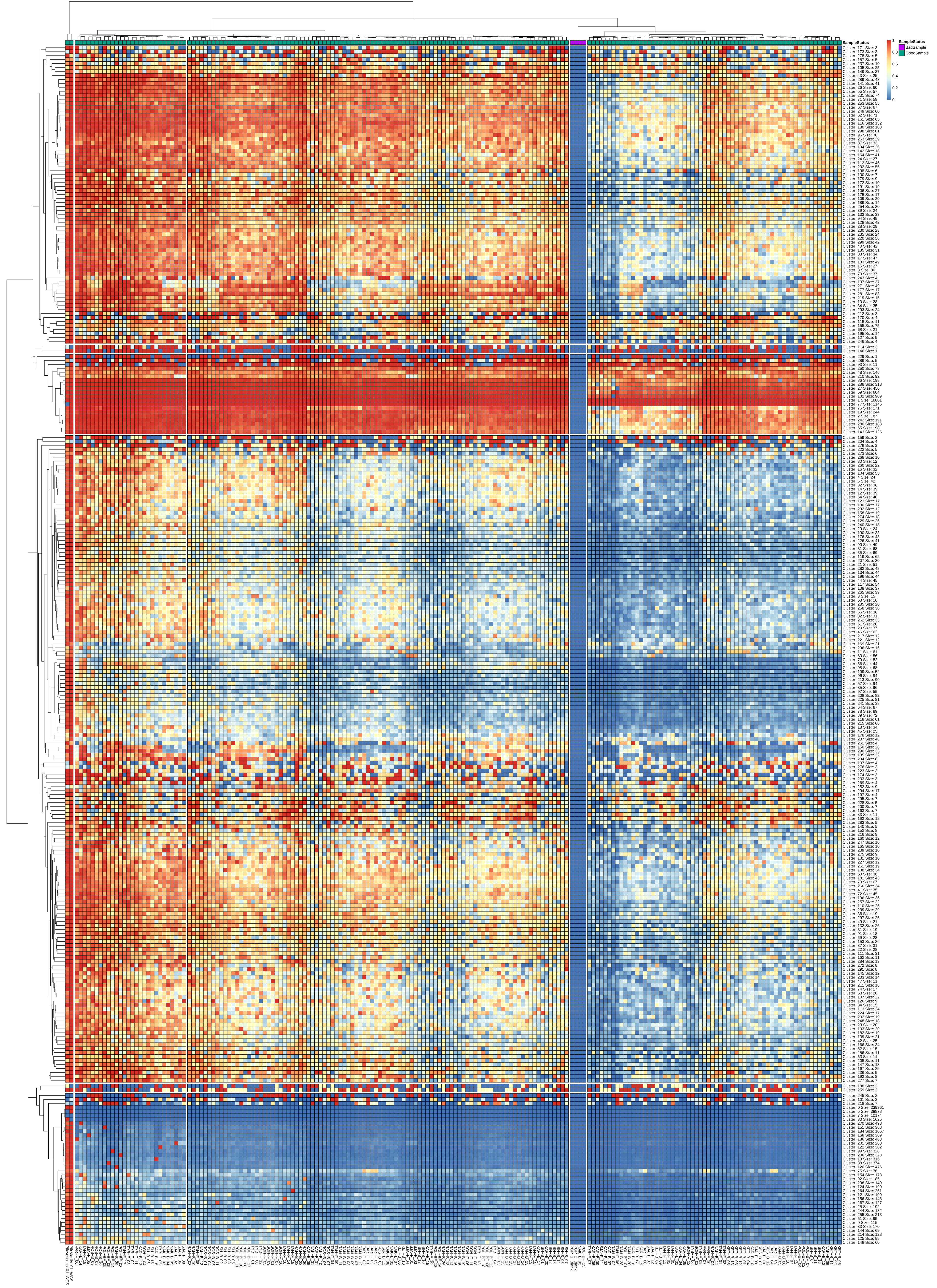

### Figure S2

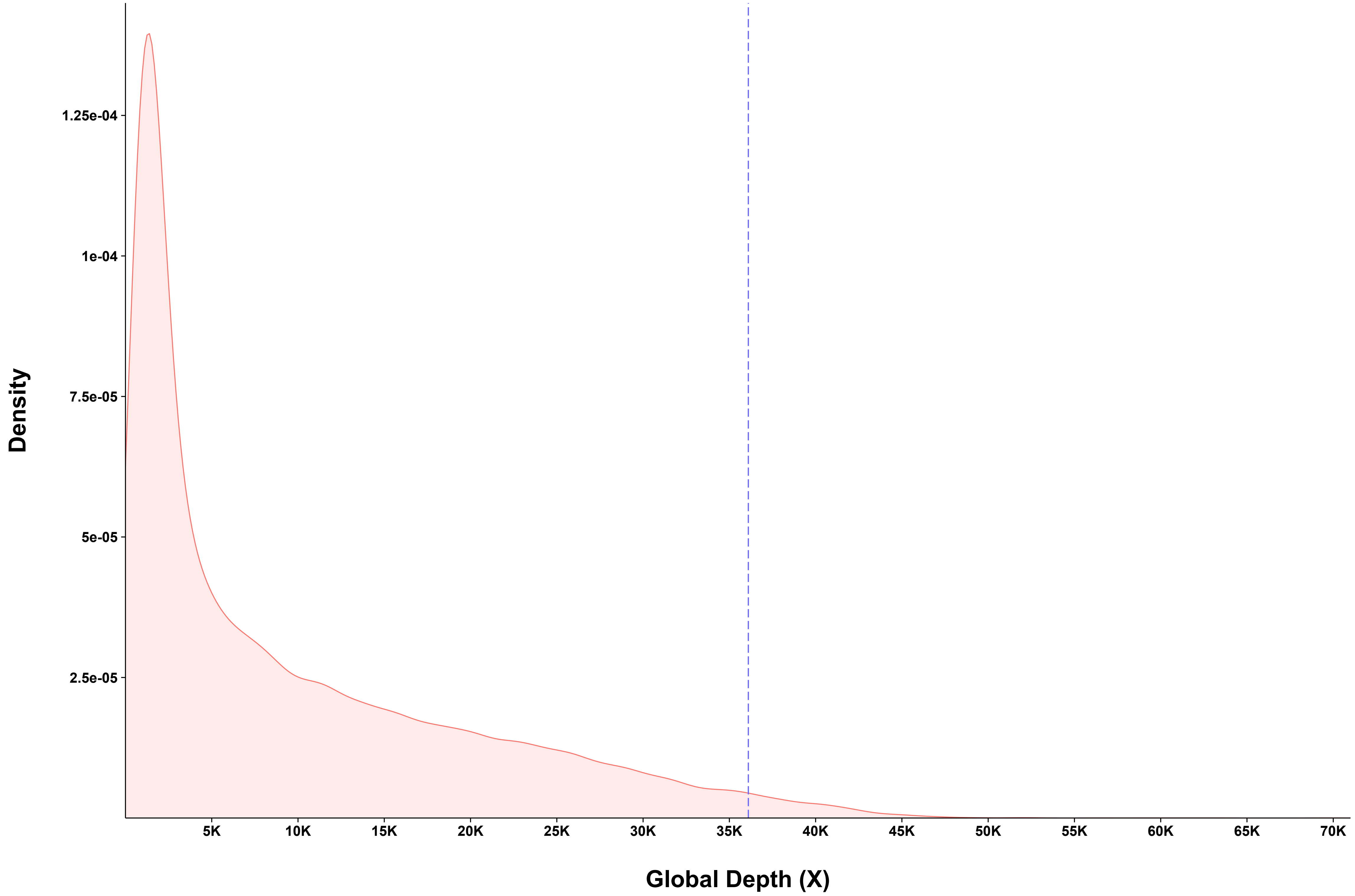
